# Aquaponics versus conventional farming: effects on the growth, nutritional and chemical compositions of *Celosia argentea* L., *Corchorus olitorius* L., and *Ocimum gratissimum* L.

**DOI:** 10.1101/2022.10.06.511176

**Authors:** Gbolaga O. Olanrewaju, David D. Sarpong, Abiola O. Aremu, Elizabeth O. Ade-Ademilua

## Abstract

This study examined the practicality and sustainability of growing leafy vegetables in aquaponics instead of traditional soil-based farming systems by comparing the physiological growth patterns, nutritional compositions, and phytochemical constituents of *Celosia argentea* L., *Corchorus olitorius* L. and *Ocimum gratissimum* L. grown in aquaponics with those of other conventional systems. The results of this study indicate that the growth and accumulation of biomass by plants grown in aquaponics were similar to those obtained in unamended loamy soil but better than those of plants grown in inorganic hydroponics. However, plants grown in NPK-supplemented soil showed significantly (p<0.05) higher biomass accumulation than those grown in aquaponics. Likewise, *C. argentea*, *C. olitorius*, and *O. gratissimum* grown in aquaponics had significantly higher nutrient compositions than those grown in inorganic medium, and at the same time, similar to that of plants grown in unamended loamy soil. *C. argentea* and *C. olitorius* grown in inorganic medium had significantly higher concentrations of the observed phytochemicals than those grown in aquaponics, whereas the opposite was true for *O. gratissimum*. The three plant species were able to serve as filters for the effective maintenance of nitrogen dynamics in the constructed African catfish aquaponics, while utilizing nitrogenous waste for biomass production.

## INTRODUCTION

The human population explosion is estimated to increase the demand for global crops and livestock by approximately 50% within the next 50 years, (Fukase & Martin, 2020) and by the year 2050, an estimated global population of 9.4 - 10 billion (United Nations, 2019a) will reach Earth’s carrying capacity. Sustainably feeding this population will require bridging a 56% food gap between the current crop production rate and the production rate needed by the year 2050, a 593 million hectare land gap between the current global agricultural land demand and 2050 demand, and an 11-gigaton greenhouse gas mitigation gap between the expected 2050 agricultural emissions and the target emission level to mitigate global warming below 2 °C (Ranganathan *et al*., 2018). Global agricultural production primarily utilizes three natural resources: arable land, water, and fossil fuel. Currently, 75% of agricultural land is degraded by human-induced erosion, salinization, compaction, nutrient depletion, and pollution, coupled with the loss of arable land to desertification at a rate of 12 million hectare per year (United Nations, 2019b, Ferreira *et al*., 2022). Over the last 20 years, nitrogen use in chemical fertilizers has exceeded the oceanic nitrogen content tremendously, (Hutchins & Capone, 2022) which has caused severe eutrophication of water bodies. Hence, the reduced availability of agricultural land and water poses a major challenge to sustainable crop production. Closing the loop between crop and animal agriculture is an effective way to improve water-use efficiency, reduce agricultural waste, and reduce environmental pollution. Aquaponics provides an effective way to reduce the dependence of future agriculture on the availability of arable land.

The concept of aquaponics, which integrates aquaculture and hydroponics, is gaining increased attention as an environmentally friendly bio-integrated food production system (Colt *et al*., 2022) with the primary goal of reusing nutrients released from fish waste for plant production. Recirculating aquaculture and hydroponics offers solutions for increasing intensive production and environmental sustainability. However, each of these systems has drawbacks that limit the overall efficiency and profitability of their operation. In recirculating aquaculture, water quality must be monitored consistently and wastewater discharge must occur regularly to maintain optimal water quality levels. Similarly, in hydroponic production, the uptake of nutrients by plants, as well as chemical changes that occur within the hydroponic solution, results in the occasional removal of water from the system, which must then be replaced by a fresh nutrient solution. Although less environmentally harmful than nutrient leaching from traditional agriculture, the disposal of nutrient water discharged from these systems presents certain challenges for producers as well as an overall loss of water conservation efficiency for the system (Baiyin *et al*., 2021). Hence, aquaponic systems, through the integration of aquaculture and hydroponics, resolve the challenges associated with conventional agricultural systems.

Despite the availability of several studies on the suitability of various aquaponic designs for various plant species, (Baiyin *et al*., 2021; Baßmann *et al*., 2020; Paudel, 2020) opportunities in the field remain largely unexplored, with less attention being paid to understanding how the system affects the growth and development of plant species and the endpoint nutritional values and phytochemical composition of plants grown in the system. Several studies have pointed out the economic benefits of aquaponic systems for leafy vegetable cultivation (Bailey & Ferrarezi, 2017; Rizal *et al*., 2018), but only a few have addressed the physiological effects of aquaponics on plants. Therefore, in this study, we comparatively assessed the effects of being grown in aquaponic cultivation on vegetative growth, biomass yield, nutritional value, and phytochemical composition of three plant species, *Celosia argentea, Corchorus olitorius*, and *Ocimum gratissimum*, which are leafy vegetables globally consumed for food and medicinal purposes (Guzzetti *et al*., 2021; Nganteng *et al*., 2022; Velsankar *et al*., 2022). By doing this, this study will provide novel information on how the three plant species are adapted to the aquaponic environment and how they can be compared to cultivation in other growth media.

## MATERIALS AND METHOD

### Seed collection and planting

Wild-type seeds of *Celosia argentea* (plumed cockscomb), *Ocimum gratissimum* (clove basil), and *Corchorus olitorius* (jute mallow) were purchased from the nursery unit of Lagos State Ministry of Agriculture, raised in the nursery for three weeks after which they were separately transplanted into various growth media. Wastewater from the aquaponics fish tank was released into the vegetable growth beds twice a week, while simultaneously draining into the aquaponic water reservoir (Figure 1). MaxiGrow pre-made hydroponics solution, used in the hydroponics, contained N (170ppm), P (31ppm), K (210ppm), Ca (90ppm), Mg (24ppm), S(32ppm), Fe (1ppm), Mn (0.25ppm), Zn (0.14ppm), B (0.16ppm), Cu (0.023ppm) and Mo (0.018ppm). The hydroponics solution concentration was adjusted as the plants grow. The pH of the hydroponics solution and the aquaponics were maintained at 5.6-6.8, while the electrical conductivity (EC) was about 1.5 mS/cm and 1.7 mS/cm respectively. The unamended soil had 18% organic carbon content (SOC), CEC of 1.35mmolkg-1, total nitrogen of 1.8gkg-1, 60.8mg of phosphorus, and 132mg of potassium. The NPK-supplemented soil contained a 14-8-20 NPK fertilizer. The amended soil had a 16.4% SOC, CEC of 3.23 mmolkg-1, total nitrogen of 2.49g/kg, phosphorus of 120.5mg, and potassium of 248mg. Both soil pH were maintained at 5.6-5.8.

**Figure 1:**
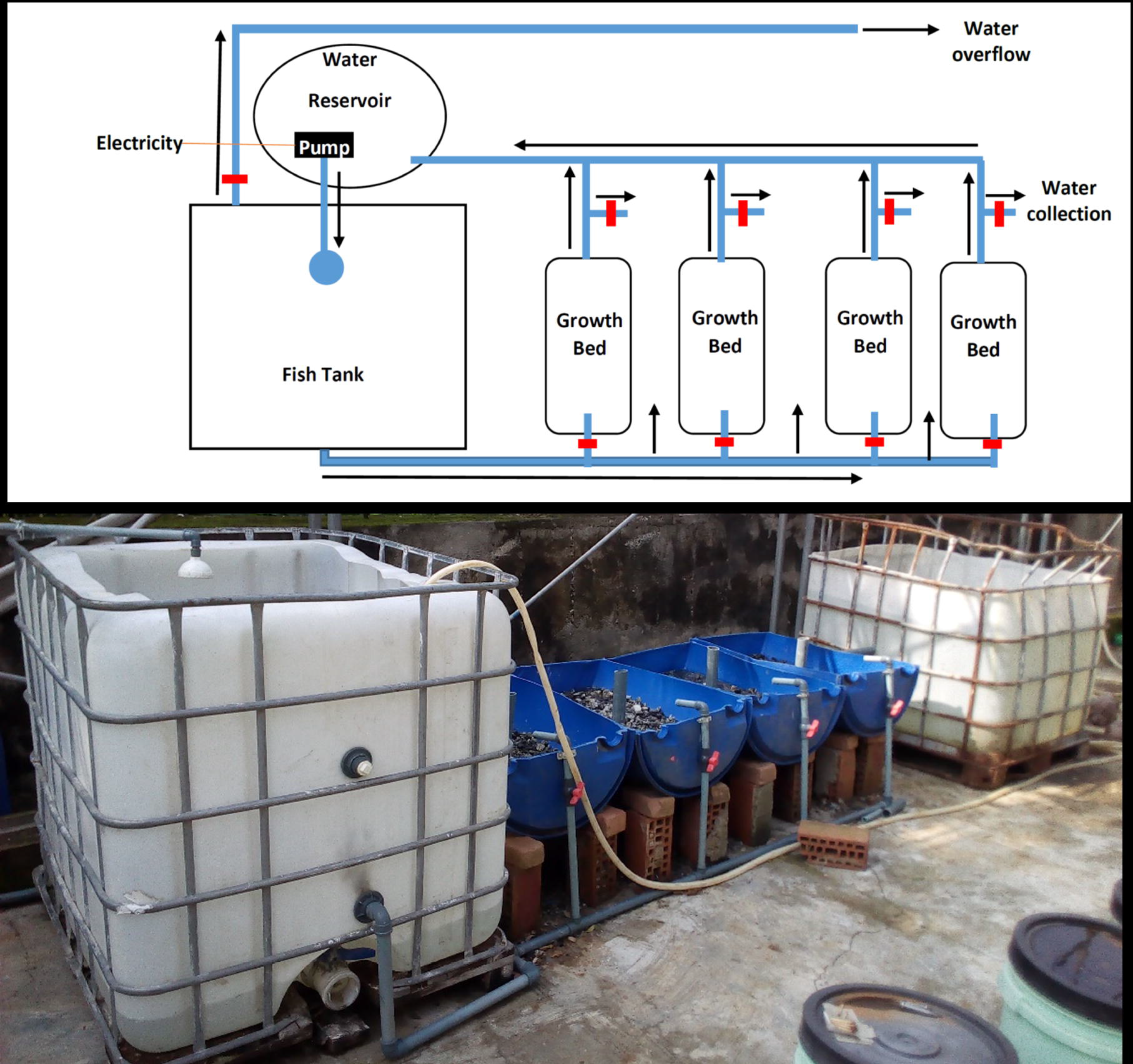
Constructed flood and drain aquaponics system. Each of the compartments contains a water valve (in red) which controls the flow of water in and out of the compartment.

### Plant growth parameters

Whole plant fresh weights were measured, after which the plant samples were bagged in an envelope and oven-dried at 65 °C for 72 h until a constant weight was achieved and the dry weight was measured. The plant heights and leaf areas were estimated using ImageJ software. All measurements were performed in triplicate and the mean values were plotted.

### Nutritional composition and phytochemical analysis

The nutritional composition of the three plant species was determined according to the AOAC protocol (Association of Official Analytical, 2005). The Sliminess test (viscosity test) of *Corchorus olitorius* was conducted according to the method described by Fasogbon and Taiwo (2019). However, viscosity was estimated as volume per time instead of area per time.

Qualitative and quantitative phytochemical analyses were performed on the methanolic extracts of the 3 plant species to determine their phytochemical constituents, as described by Kulshreshtha and Saxena (2022).

### Ammonium, nitrite and nitrate concentrations

The concentrations of non-ionized ammonia, nitrite, and nitrate in the collected water samples were determined according to the methods described in Schullehner et al. (2017)

#### Statistical analysis

The mean and standard error of the experimental triplicates were computed using the R software. Statistical tests for significance were performed using analysis of variance and Student’s t-test (P < 0.05).

## RESULTS

### Fresh and Dry weights

To assess how the constructed flood and drain aquaponics system (Figure 1) affects plant growth and development, this study measured the fresh and dry weights, number of leaves, and total leaf areas of *Celosia argentea, Corchorus olitorius*, and *Ocimum gratissimum* vegetables grown in the Aquaponics system, over 10 weeks growth duration, and compared them to those of vegetables grown in the NPK fertilized soil, inorganic hydroponic system, and unamended loamy soil. The fresh and dry weights exhibited a similar distribution pattern across the three plants, with plants grown in NPK-supplemented soils having significantly higher (p<0.05) fresh and dry weights in the 10th week than in the others. The fresh and dry weights of plants grown in aquaponics and unamended loamy soil were not significantly different from each other but were significantly higher (p<0.05) than those grown in an inorganic hydroponic system and tap water in the 10th week (Figure 2). Over the 10 weeks of observation, *C. argentea* cultivated in the Aquaponics system maintained fresh weights that were not significantly different from those grown in unamended loamy soil and inorganic hydroponics but were significantly lower than the fresh weight of *C. argentea* grown in NPK-supplemented soil. A slight difference was observed in the dry weight of *C. argentea* at the 10th week, with those grown in the aquaponic system having a significantly higher dry weight than those grown in inorganic hydroponic medium (Figure 2a).

**Figure 2:**
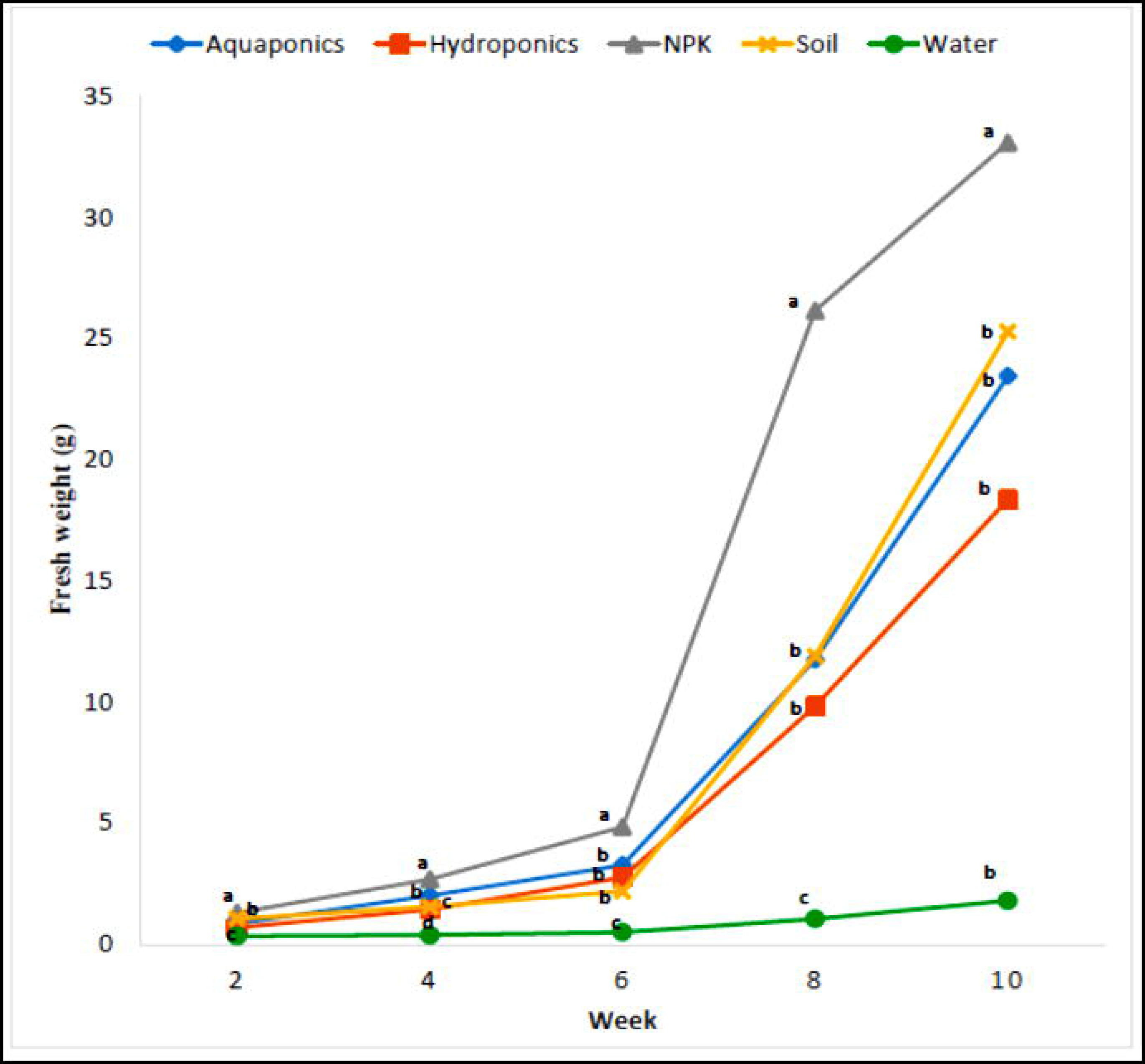
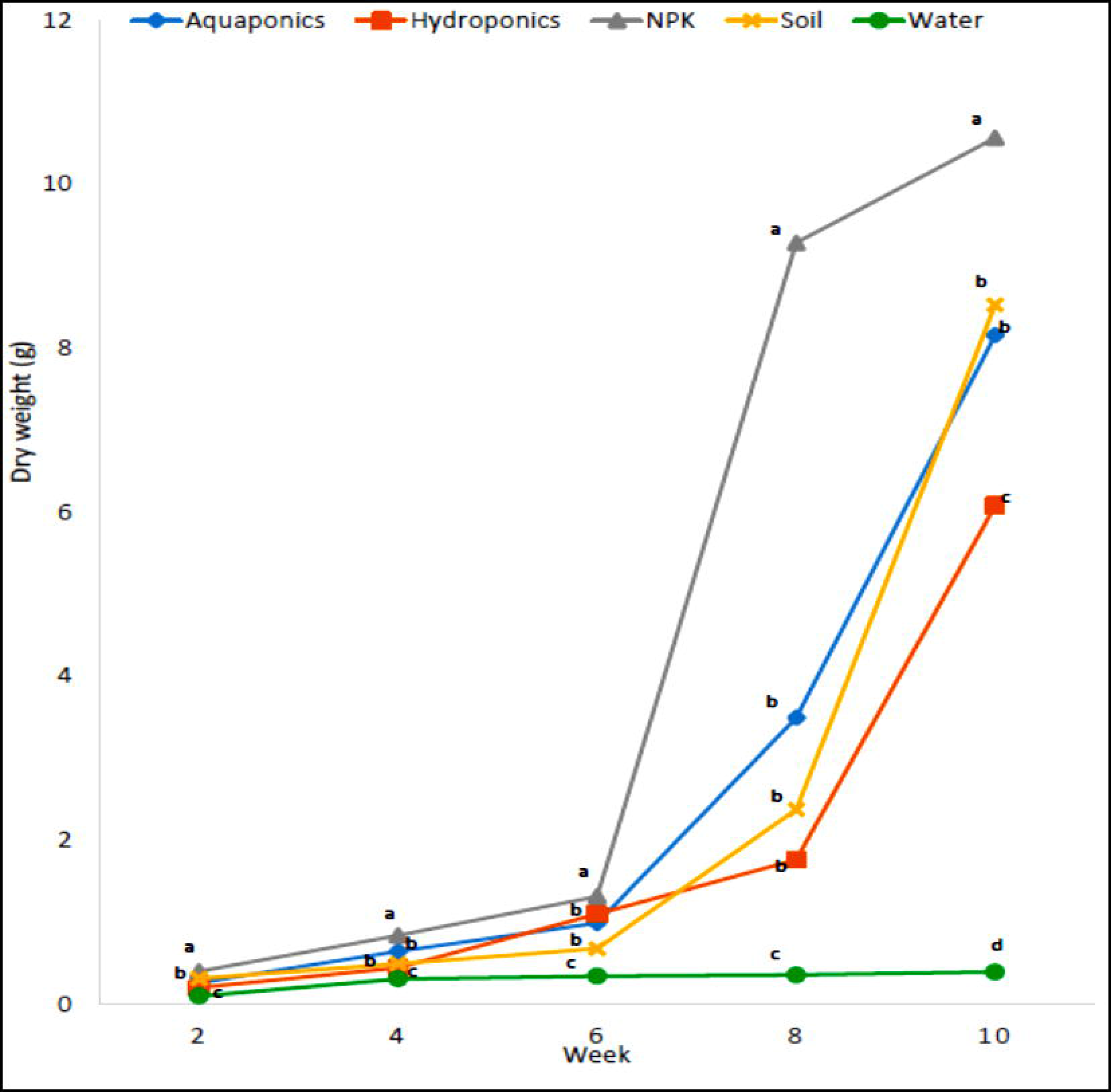
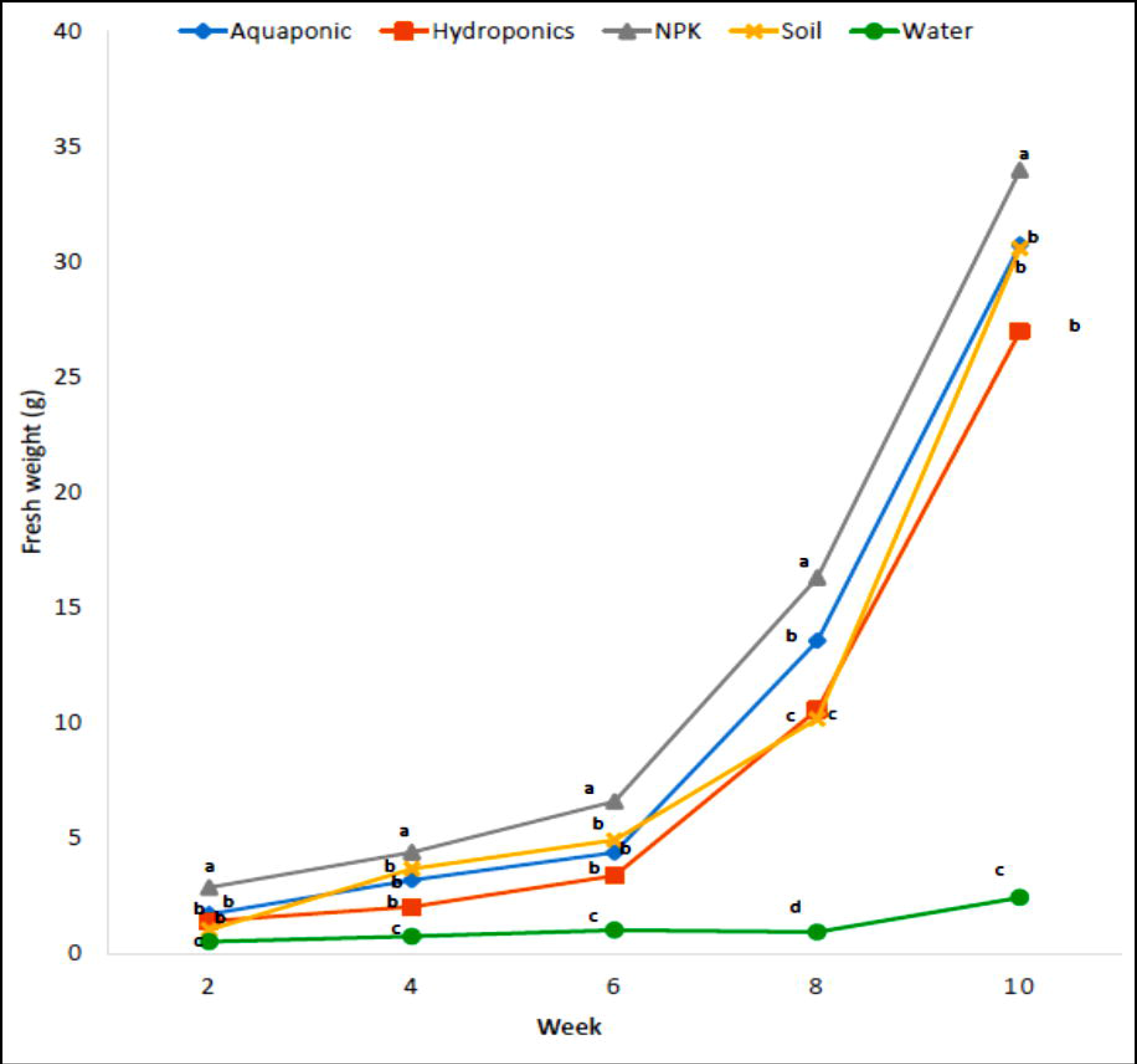
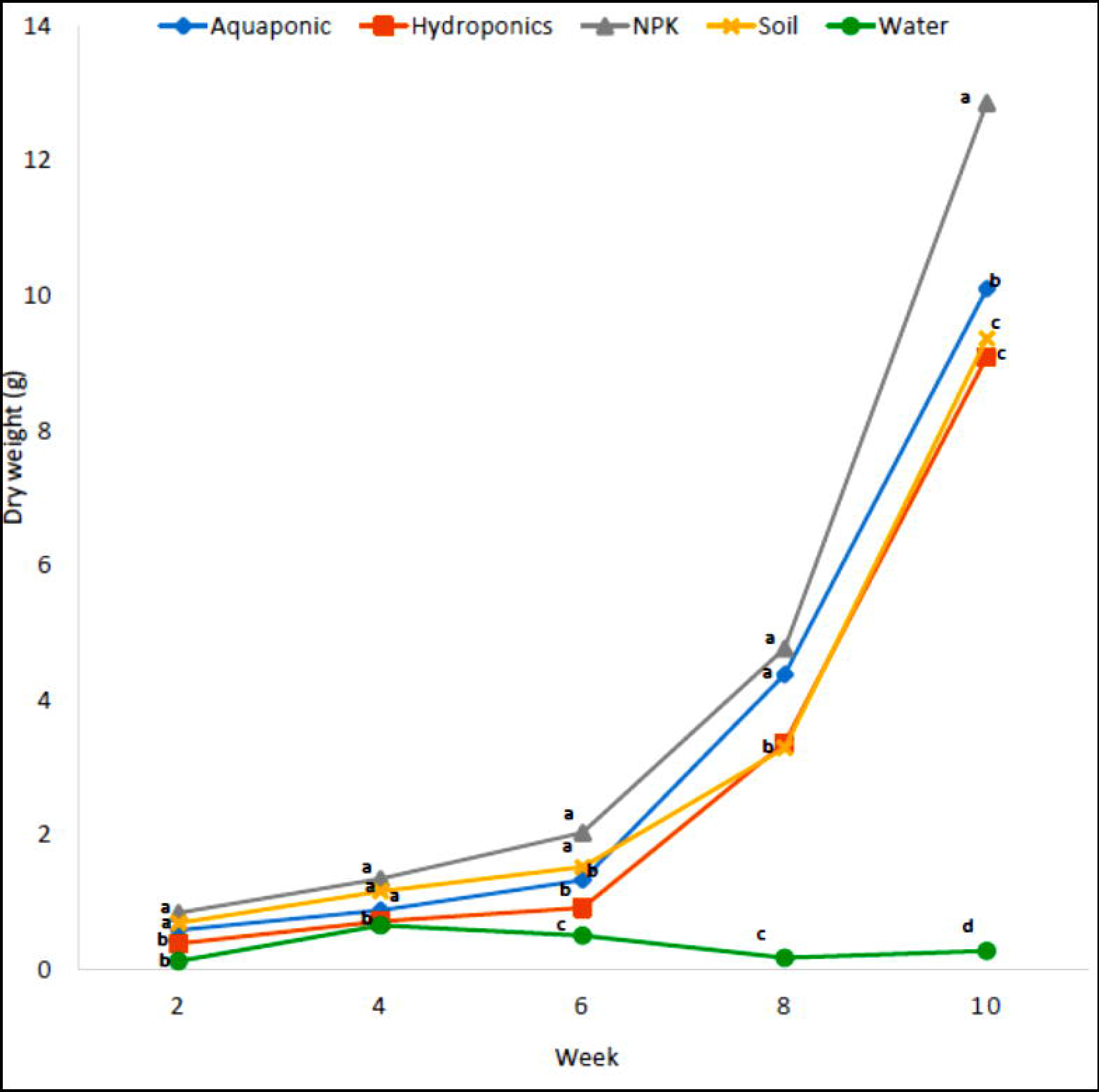
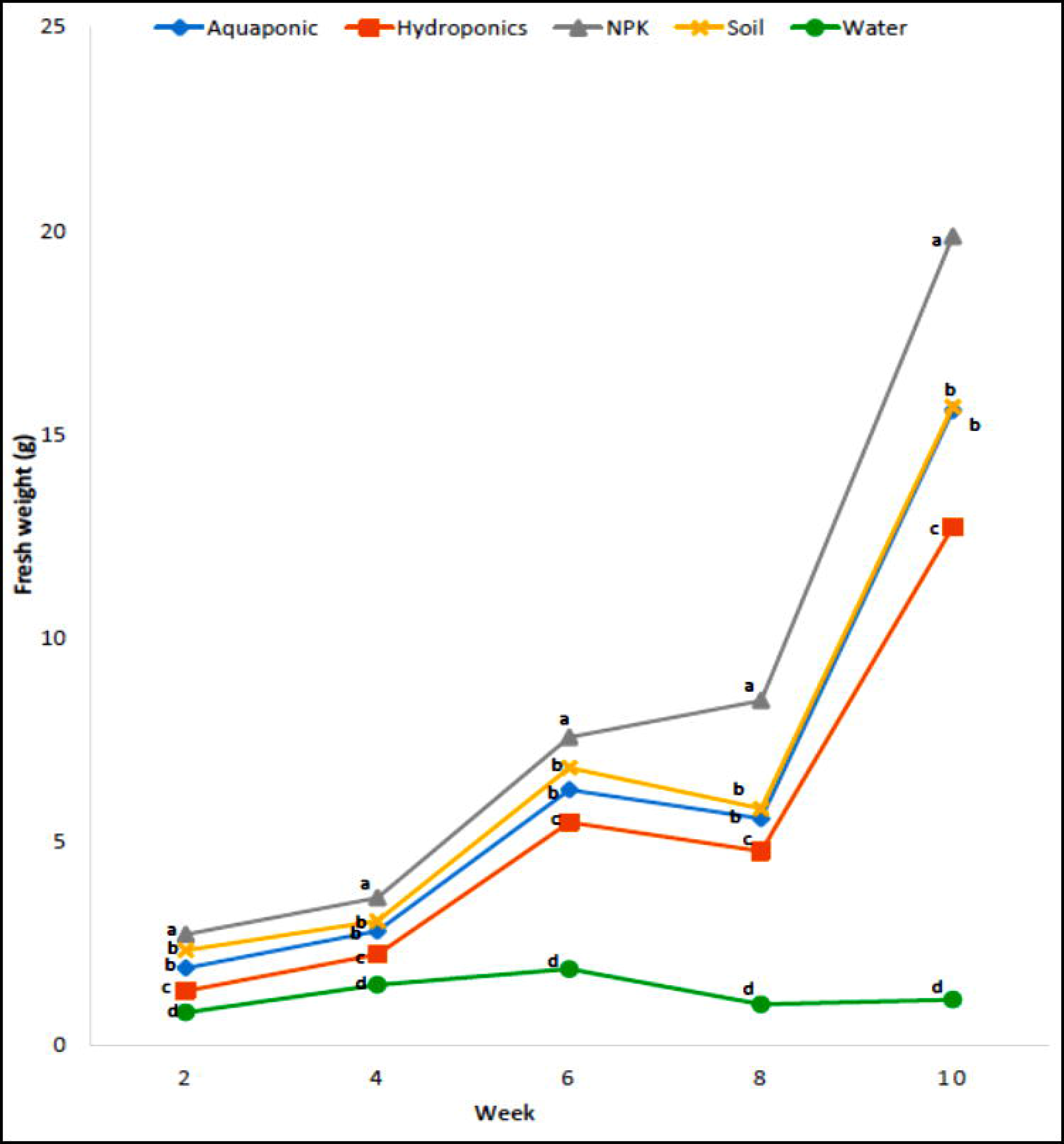
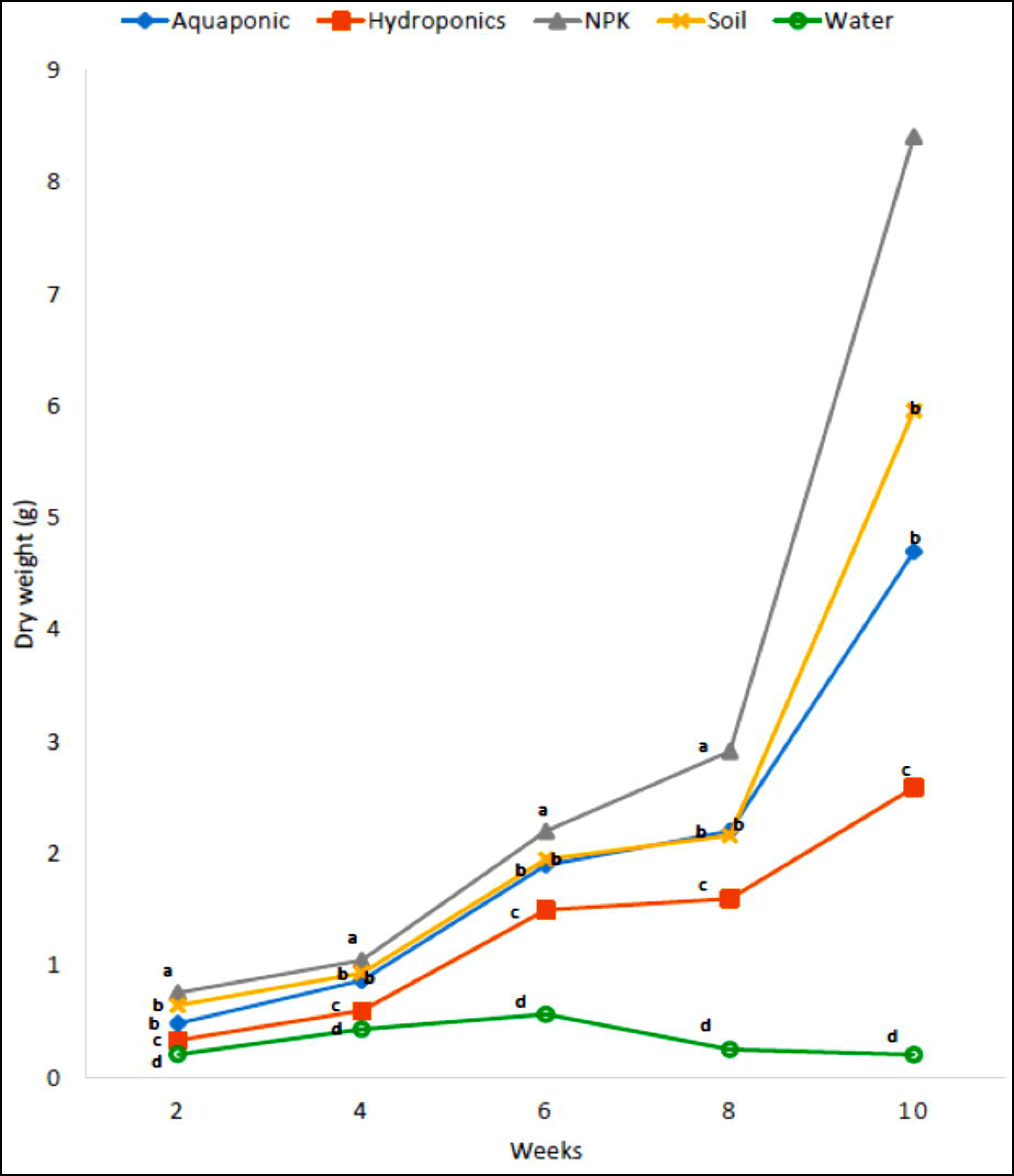
Fresh and dry weights of the plant species grown in the different growth medium. The fresh weights are in the upper panel while the dry weights are in the middle panels. **a**. *Celosia argentea* **b**. *Corchorus olitorius* **c**. *Ocimum gratissimum*. Aquaponics (blue), inorganic hydroponics (orange), NPK supplemented soil (grey), unamended loamy soil (yellow) and tap water (green). Plotted means, in the same week and on the same horizontal axis, represented with different letters are significantly different at p= 0.05. On the lower panel is the image of the 3 plant species harvested on the 10th week. Leftmost is *C. argentea, the* middle is *C. olitorius* and the far right is *O. gratissimum*. A = aquaponics, N= NPK supplemented soil, H= hydroponics and L = unamended loamy soil.

Similarly, *C. olitorius* grown in aquaponics maintained a fresh weight that was consistent with that of *C. olitorius* grown in unamended loamy soil and inorganic hydroponics (Figure 2b). However, the dry weight of *C. olitorius* grown in the aquaponic system was significantly higher than that of the unamended loamy soil and inorganic hydroponics. *C. olitorius* grown in the NPK-amended soil consistently maintained significantly higher fresh and dry weights over the 10 weeks of observation (Figure 2b). The fresh weight of *O. gratissimum* grown in the aquaponic system was significantly similar to that of *O. gratissimum* grown in the unamended loamy soil, and both were significantly greater than that of *O. gratissimum* grown in the inorganic hydroponic system. A similar pattern of weight accumulation was observed in the dry weight of *O. gratissimum*, with the NPK-supplemented soil producing *O. gratissimum* with consistently significantly higher fresh and dry weights throughout the 10 weeks of observation (Figure 2c). The distribution of fresh and dry weights across the three plant species was highlighted as follows: **Fresh weight**: NPK > aquaponics/loamy soil/inorganic phosphate > tap water (α =0.05) and **Dry weight**: NPK > aquaponics/loamy soil > inorganic phosphate > tap water (α =0.05).

### Plant heights

The differences in height between *Celosia argentea* grown in aquaponics and those grown in the unamended loamy soil were not significant, but they were both significantly taller than those grown in inorganic hydroponic and tap water (Figure 3a). *C. argentea* grown in NPK-supplemented soil were significantly taller than the others at all weeks of observation, excluding the 8th week. With the exception of the 2nd week of observation, when no significant difference (p> 0.05) in height was observed, *Ocimum gratissimum* grown in NPK-supplemented soil were significantly taller (p< 0.05) than *O. gratissimum* grown in aquaponics throughout the observation period (Figure 3b). *O. gratissimum* grown in aquaponics maintained significantly greater heights (p< 0.05) than *those* grown in inorganic hydroponics and tap water throughout the observation period. A comparison of the heights of *O. gratissimum* grown in unamended loamy soil and aquaponics showed no significant differences (p> 0.05) over the weeks of analysis. However, there was no significant difference in the height of *Corchorus olitorius* grown in aquaponics and those grown in NPK supplements soil. *C. olitorius* grown in these two media were significantly taller (p<0.05) than those grown in the other growth media (Figure 3c).

**Figure 3:**
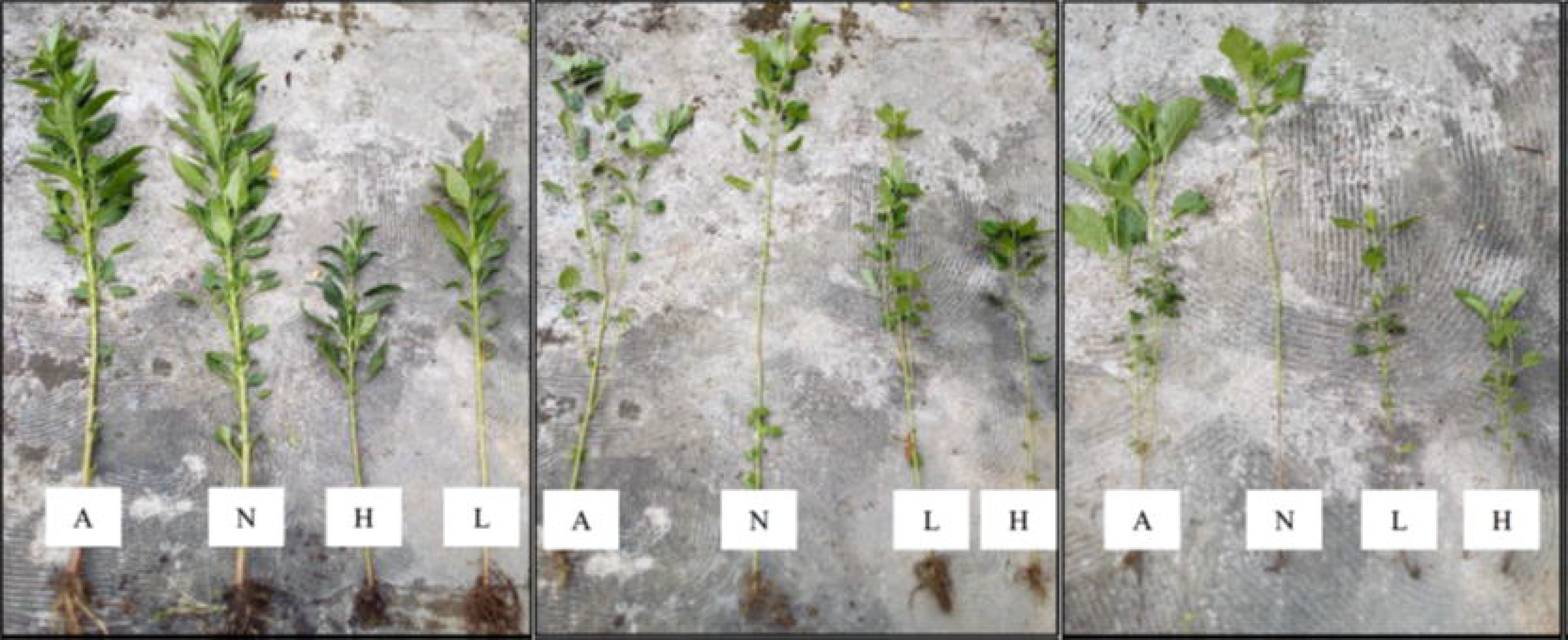
Heights of the plant species grown in the different growth medium. **a**. *Celosia argentea* **b**. *Ocimum gratissimum* **c**. *Corchorus olitorius*. Aquaponics (blue), inorganic hydroponics (orange), NPK supplemented soil (grey), unamended loamy soil (yellow) and tap water (green). Plotted means, in the same week and on the same horizontal axis, represented with different letters are significantly different at p= 0.05.

### Total leaf areas and number of leaves

*Celosia argentea* grown in aquaponics had leaves with significantly larger (p< 0.05) total area than those grown in inorganic hydroponics throughout the experimental period. At 8th and 10th weeks of growth, the total leaf surface area was not significantly different between *C. argentea* grown in aquaponics, NPK-supplemented soil, and unamended loamy soil (Figure 4a). Similarly, at the 10th week of growth, the leaves of *O. gratissimum* grown in aquaponics had surface areas that were not significantly different from those grown in NPK-supplemented soil and unamended loamy soil (Figure 4b). Unlike the other two plant species, the leaf surface areas of *Corchorus olitorius* grown in aquaponics were not significantly different from those grown in inorganic hydroponics and unamended loamy soil but were significantly lower than those grown in NPK-supplemented soil (Figure 4c).

**Figure 4:**
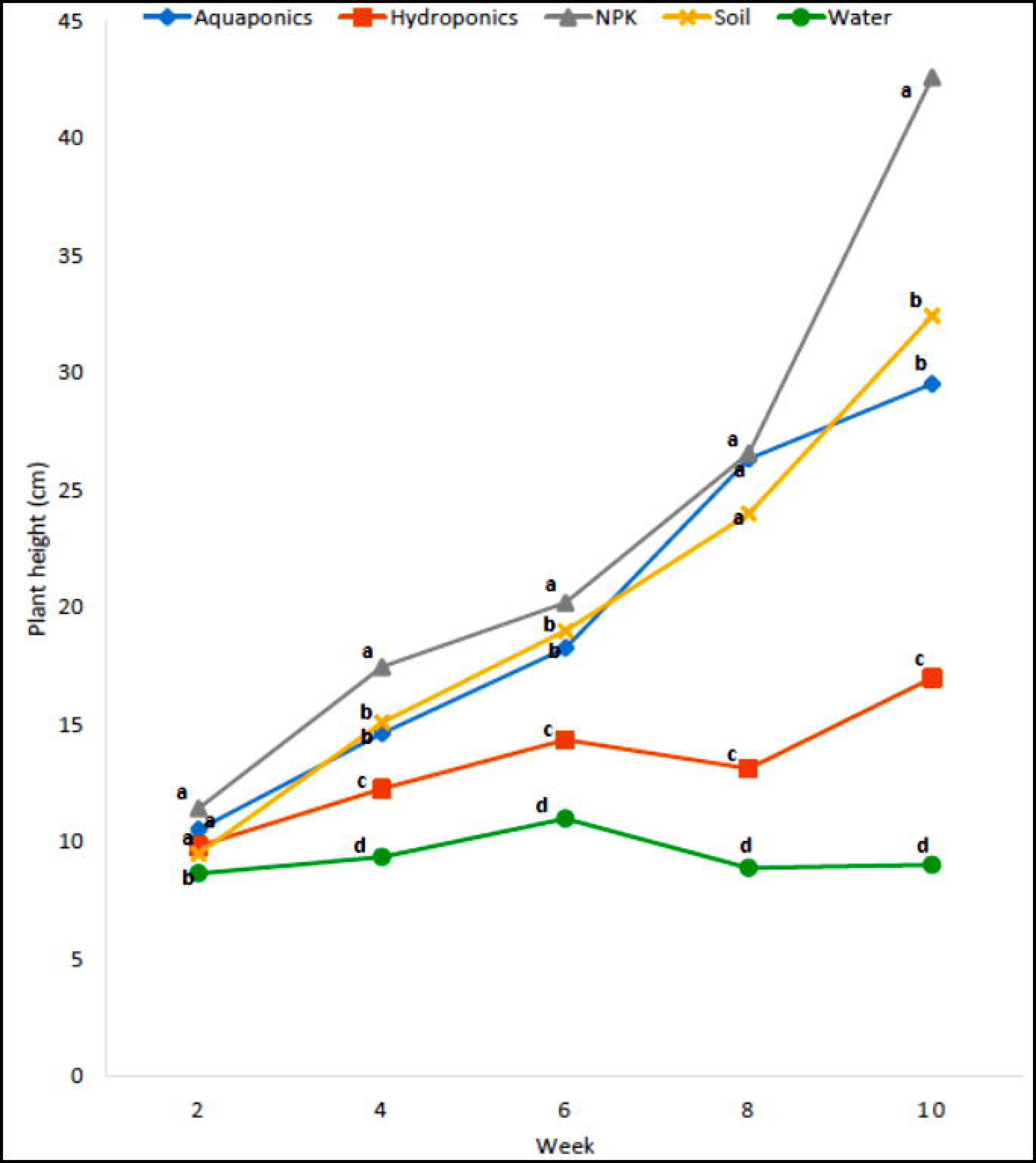
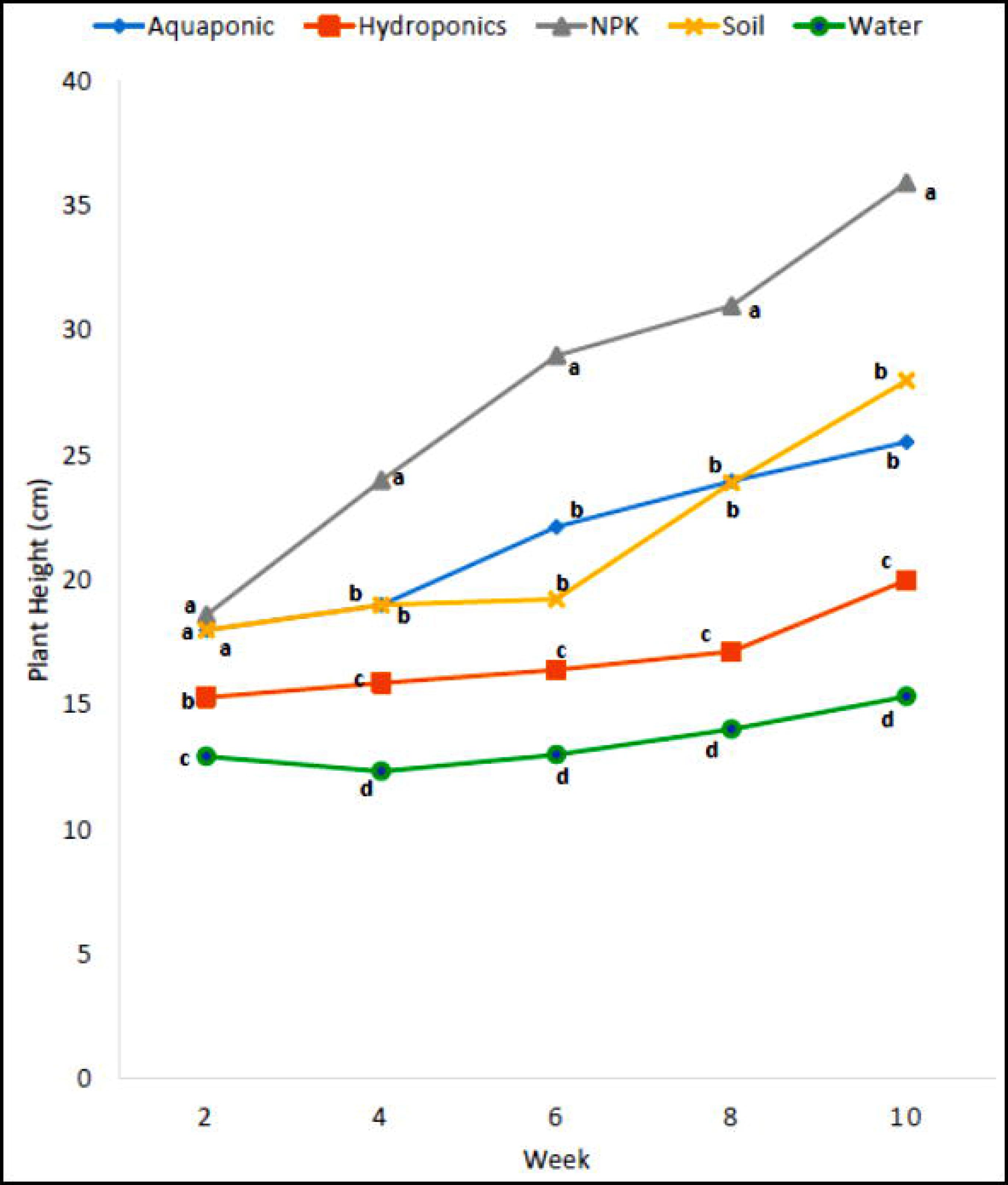
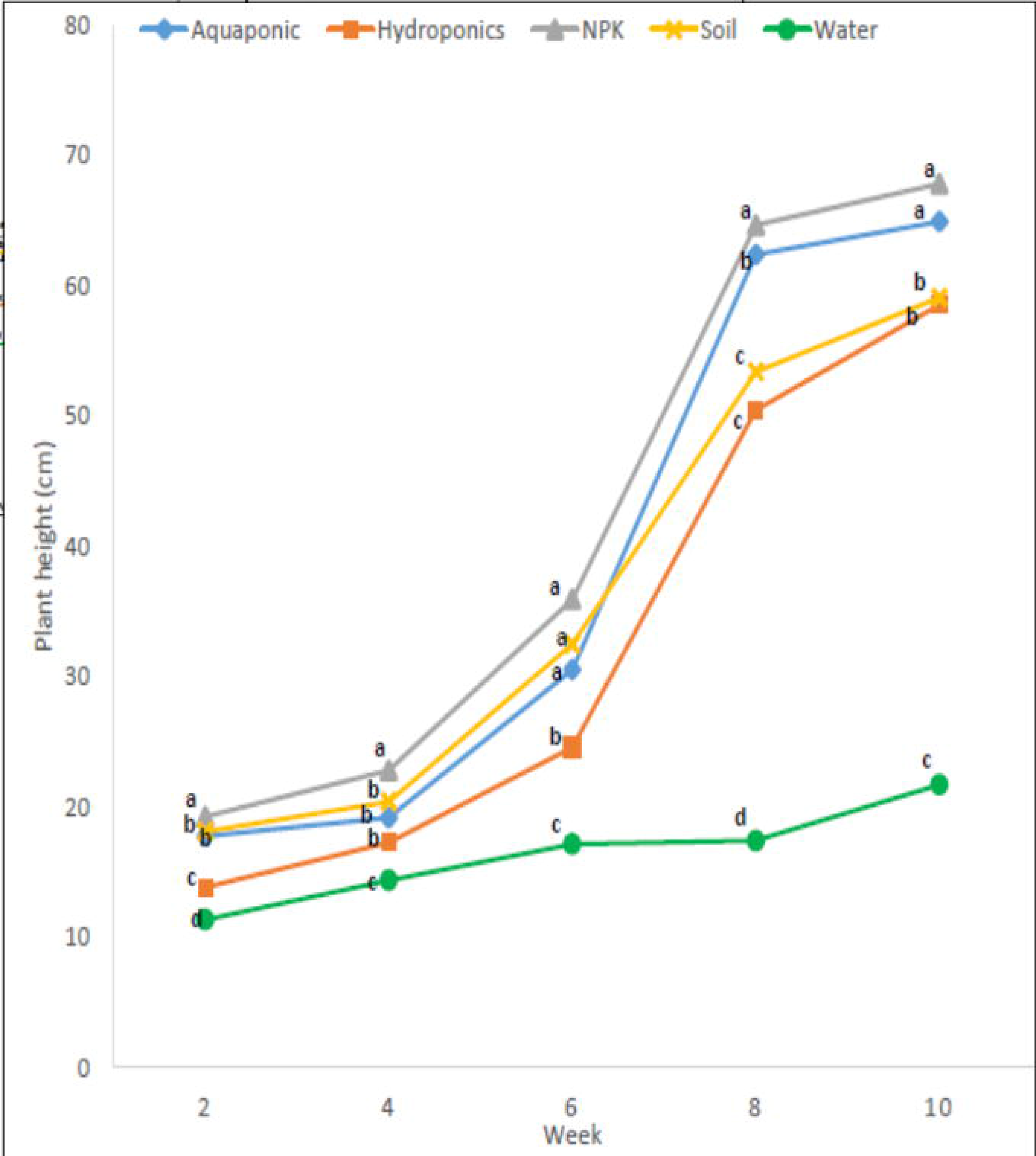
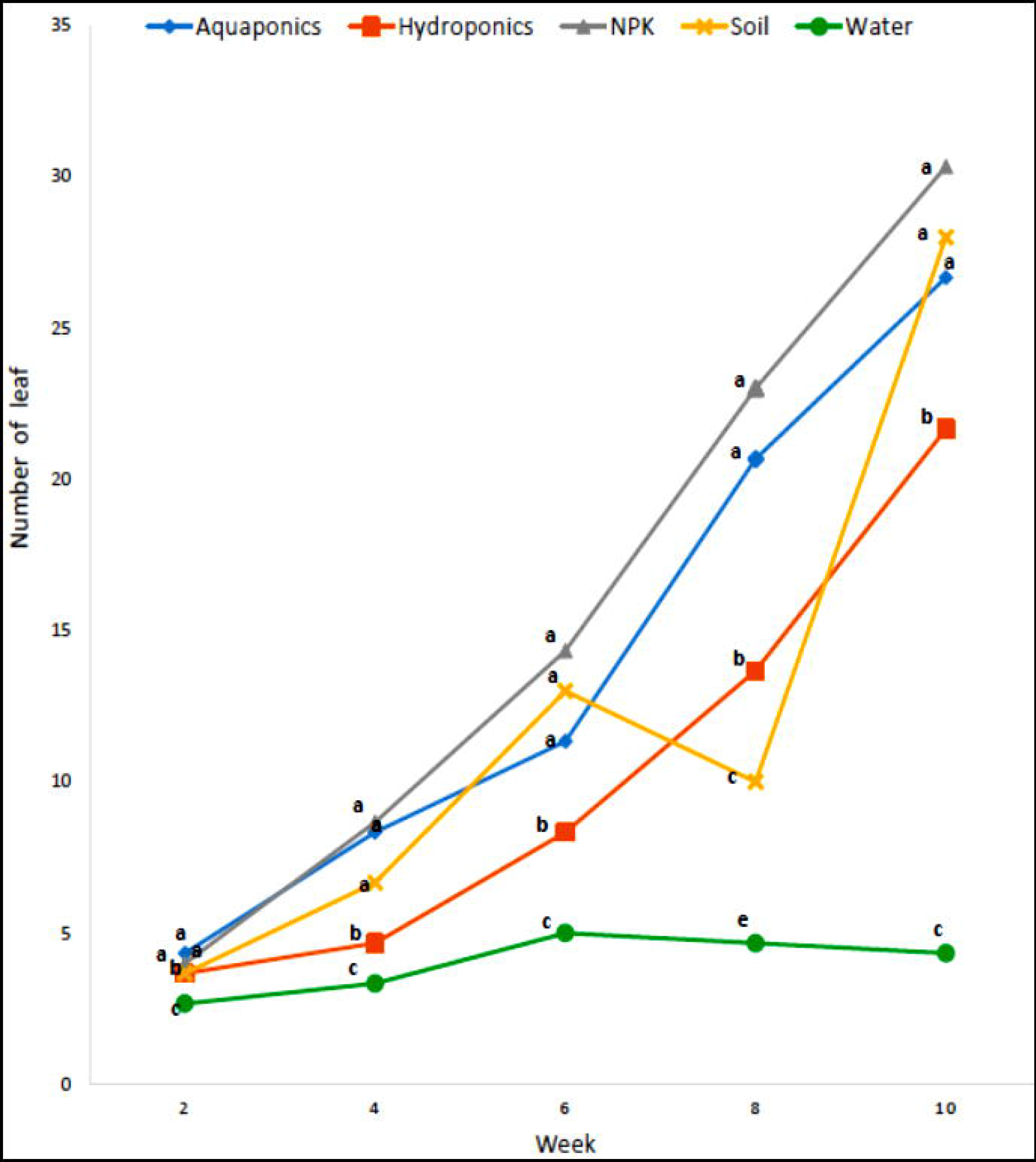
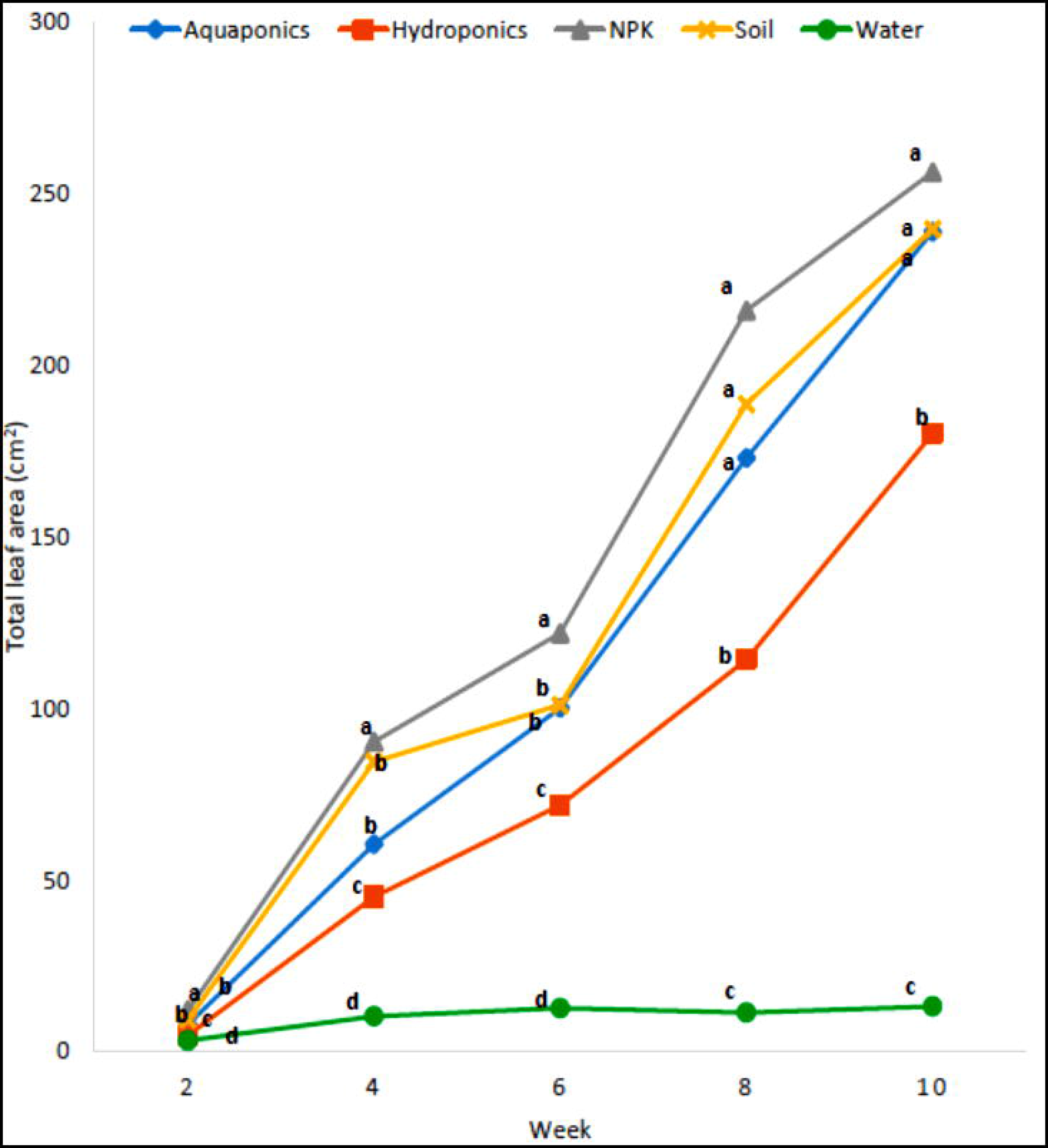
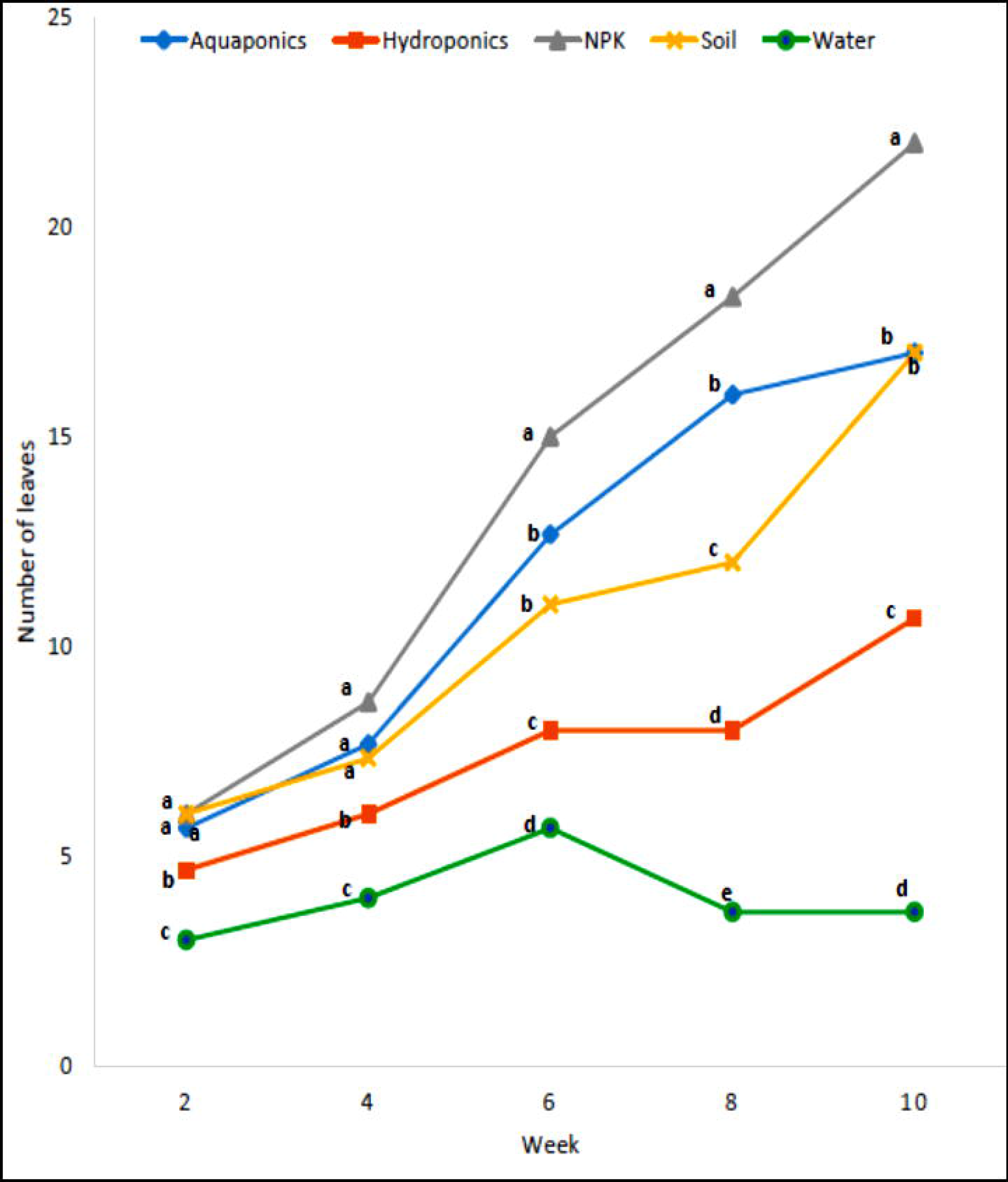
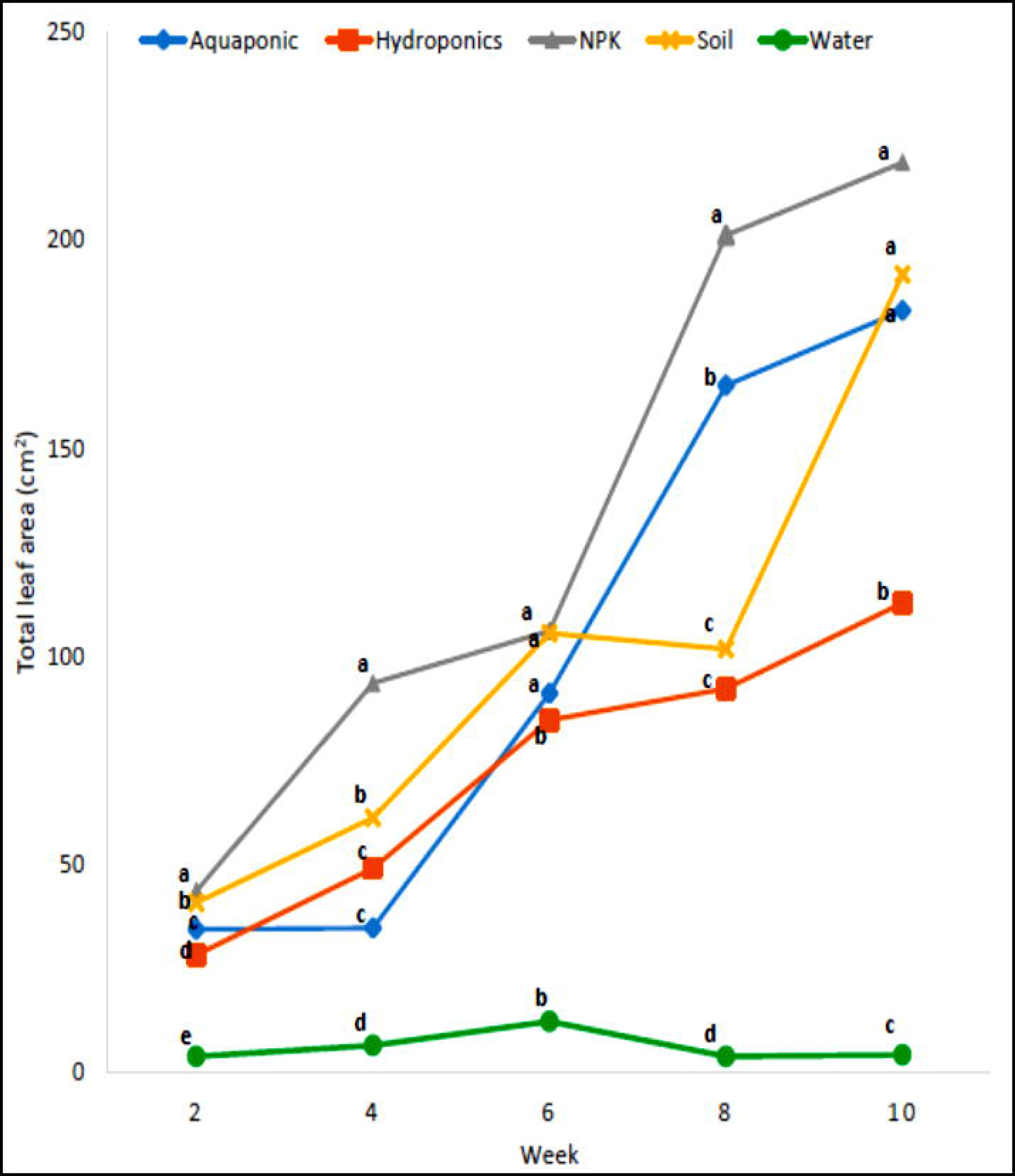
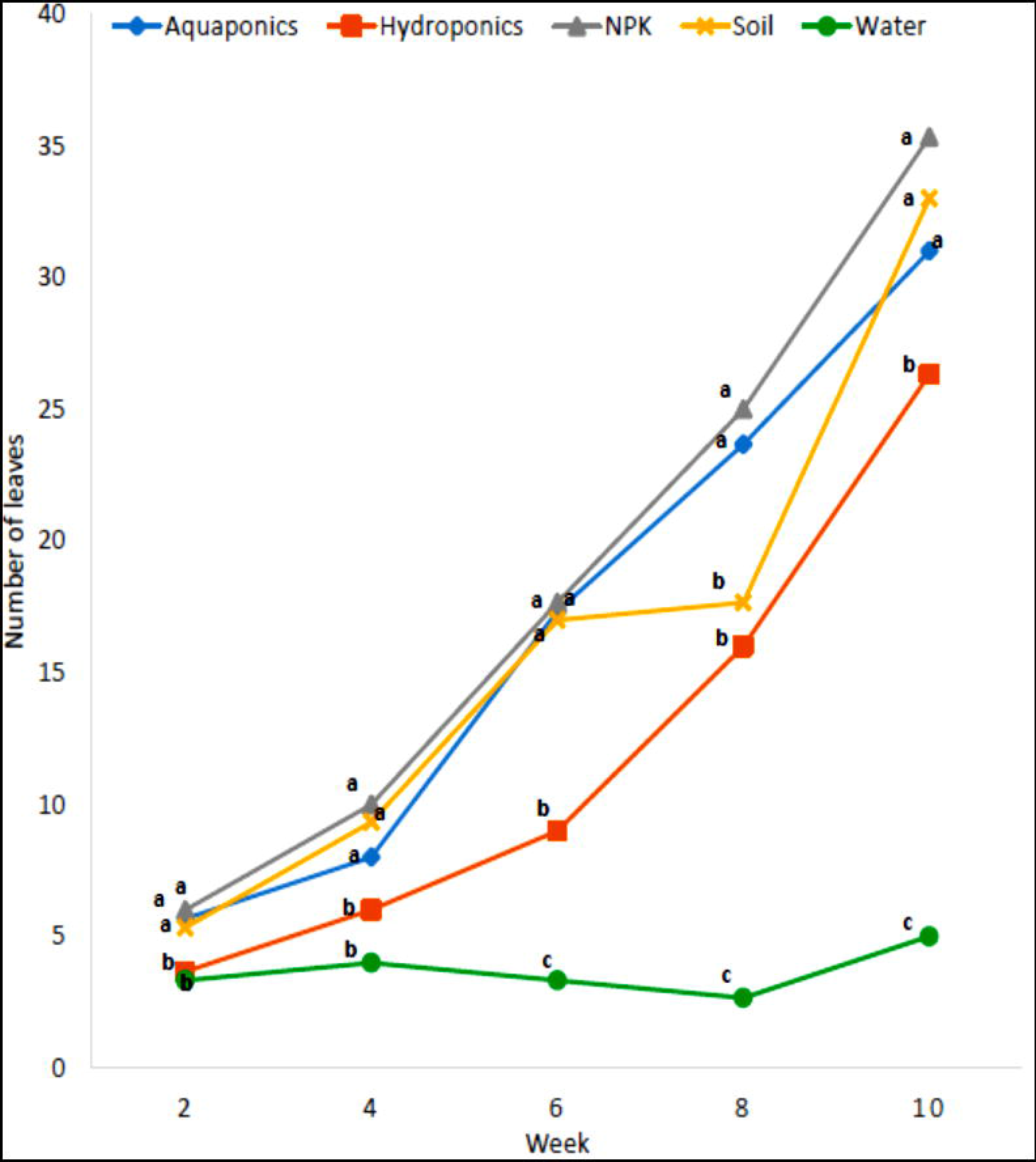
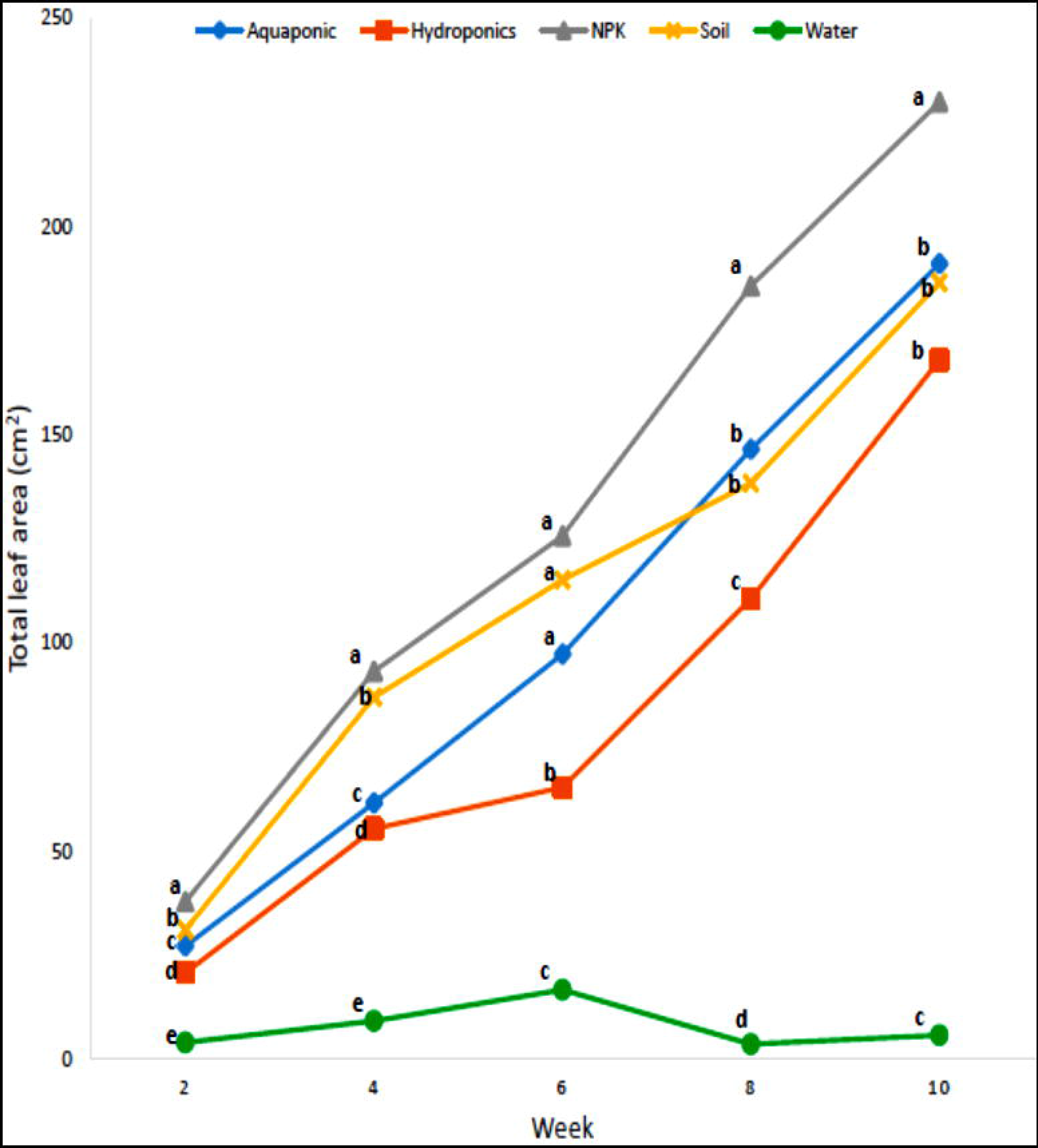
Number and total leaf areas of the plant species grown in the different growth medium. The number of leaves produced by each plant are in the upper panel while the total leaf area is in the lower panels. **a**. *Celosia argentea* **b**. *Ocimum gratissimum* **c**. *Corchorus olitorius*. Aquaponics (blue), inorganic hydroponics (orange), NPK supplemented soil (grey), unamended loamy soil (yellow) and tap water (green). Plotted means, in the same week and on the same horizontal axis, represented with different letters are significantly different at p= 0.05.

At the 10th week, the average number of leaves produced by both *C. argentea* and *C. olitorius* grown in aquaponics was not significantly different from that of plants grown in both NPK-supplemented and unamended loamy soils (Figure 4a and c). However, the average number of leaves in *O. gratissimum* grown in aquaponics was statistically similar to that of *O. gratissimum* grown in unamended loamy soil, and both were significantly lower than the average number of leaves in *O. gratissimum* grown in NPK-supplemented soil (Figure 4b). Common to all three plant species was the production of a higher number of leaves in aquaponic plants than in plants grown in inorganic hydroponics. There was a disruption in the number of leaves between the 6th and 8th month in plants grown in unamended loamy soils (Figure 4).

### Nutritional analysis

Plants are commonly consumed for their nutritional and medicinal benefits (Ahnen *et al*., 2019), which are often affected by their growth conditions and environment (Chrysargyris *et al*., 2020). Hence, to assess the effects of the aquaponic growth medium on the nutritional and phytochemical composition of the three plant species, we conducted a proximate analysis of the 10weeks old leaves to identify the nutritional composition of plant matter, a phytochemical analysis to identify the common metabolites of medicinal interest, and a sliminess test on *C. olitorius* to identify its palatability as part of a Nigerian cuisine. The leaves of *Celosia argentea* grown in aquaponics had a significantly higher carbohydrate content than those of *C. argentea* grown in NPK-supplemented soil and inorganic hydroponics. Carbohydrate content in the leaves of *C. argentea* grown in aquaponics was not significantly different from that in the unamended loamy soil (Table 1a). A similar trend was observed in protein content, with aquaponics grown *in C. argentea* having a similar protein content to those grown in unamended loamy soil, and both had a significantly higher protein content than *C. argentea* grown in inorganic hydroponics and NPK-supplemented soil. A reverse situation was observed in the crude fat and ash contents, as *C. argentea* grown in aquaponics and those grown in the unamended loamy soil had significantly lower crude fat and ash contents than the rest (Table 1a). Moisture and crude fiber contents were not significantly affected by the aquaponic system.

**Table 1a:** Nutritional constituent of *Celosia argentea* grown in the four growth medium. Values with different letters are significantly different across the row at α level of 0.05.

| Parameters (%) | Aquaponics | Hydroponics | NPK soil | Loamy Soil |
| --- | --- | --- | --- | --- |
| CHO | 23.99 + 1.34<sup>a</sup> | 21.70 + 1.18<sup>b</sup> | 19.39 + 1.13<sup>b</sup> | 24.49 + 1.23<sup>a</sup> |
| Protein | 6.65 + 0.13<sup>a</sup> | 6.25 + 0.23<sup>b</sup> | 6.15 + 0.21<sup>c</sup> | 6.85 + 0.12<sup>a</sup> |
| Crude fat | 2.49 + 0.21<sup>b</sup> | 3.09 + 0.18<sup>a</sup> | 3.39 + 0.14<sup>a</sup> | 2.99 + 0.22<sup>b</sup> |
| Moisture | 35.88 + 2.12<sup>a</sup> | 36.5 + 2.04<sup>a</sup> | 36.78 + 2.06<sup>a</sup> | 36.33 + 1.91<sup>a</sup> |
| Ash | 1.97 + 0.09<sup>c</sup> | 2.58 + 0.2<sup>b</sup> | 3.125 + 0.18<sup>a</sup> | 2.04 + 0.18<sup>c</sup> |
| Crude fibre | 29.05 + 1.13<sup>a</sup> | 30.6 + 1.24<sup>a</sup> | 31.55 + 1.21<sup>a</sup> | 29.55 + 1.12<sup>a</sup> |

*Corchorus olitorius* grown in the aquaponics system had carbohydrate, protein, and crude fat contents that were significantly higher than those of *Corchorus olitorius* grown in NPK-supplemented soil and inorganic hydroponics, and were not significantly different from those of *Corchorus olitorius* grown in the unamended loamy soil (Table 1b). *Corchorus olitorius* grown in aquaponics also retained moisture content that was not significantly different from that grown in other growth media, with the exception of NPK-supplemented soil, which had a significantly higher moisture content. *Corchorus olitorius* grown in aquaponics also had the lowest ash and crude fiber contents among the four growth media (Table 1b). The nutritional composition of *Ocimum gratissimum* also followed a similar pattern to those grown in aquaponics and unamended loamy soil, with significantly higher carbohydrate, protein, and ash content than those grown in other growth media. Aquaponics grown *in O. gratissimum* had a significantly higher crude fiber content than those grown in inorganic hydroponics and NPK-supplemented soil, while the moisture and crude fat content was at par with the other growth media, except for the NPK-supplemented soil and inorganic hydroponics, respectively (Table 1c).

**Table 1b:** Nutritional constituent of *Corchorus olitorius* grown in the four growth medium Values with different letters are significantly different across the row at α level of 0.05.

| Parameters (%) | Aquaponics | Hydroponics | NPK | Soil |
| --- | --- | --- | --- | --- |
| CHO | 24.91 + 1.04<sup>a</sup> | 23.6 + 1.08<sup>b</sup> | 22.34 + 1.09<sup>b</sup> | 25.32 + 1.14<sup>a</sup> |
| Protein | 7.39 + 0.14<sup>a</sup> | 6.92 + 0.23<sup>b</sup> | 6.85 + 0.17<sup>b</sup> | 7.59 + 0.21<sup>a</sup> |
| Crude fat | 8.17 + 0.11<sup>a</sup> | 7.76 + 0.14<sup>b</sup> | 6.38 + 0.14<sup>c</sup> | 7.97 + 0.24<sup>a</sup> |
| Moisture | 41.48 + 2.31<sup>b</sup> | 42.8 + 2.16<sup>b</sup> | 48.6 + 2.03<sup>a</sup> | 42.9 + 1.64<sup>b</sup> |
| Ash | 2.16 + 0.10<sup>c</sup> | 2.70 + 0.09<sup>b</sup> | 2.85 + 0.18<sup>b</sup> | 3.76 + 0.14<sup>a</sup> |
| Crude fibre | 15.86 + 1.13<sup>b</sup> | 17.40 + 1.24<sup>a</sup> | 18.32 + 1.21<sup>a</sup> | 17.46 + 1.12<sup>a</sup> |

**Table 1c:** Nutritional constituent of *Ocimum gratissimum* grown in the four growth medium. Values with different letters are significantly different across the row at α level of 0.05.

| Parameters (%) | Aquaponics | Hydroponics | NPK | Soil |
| --- | --- | --- | --- | --- |
| CHO | 26.80 + 1.18<sup>a</sup> | 23.1 + 1.03<sup>b</sup> | 23.04 + 1.01<sup>b</sup> | 27.00 + 1.16<sup>a</sup> |
| Protein | 4.72 + 0.12<sup>a</sup> | 4.41 + 0.21<sup>b</sup> | 4.45 + 0.14<sup>b</sup> | 4.70 + 0.11<sup>a</sup> |
| Crude fat | 4.07 + 0.13<sup>b</sup> | 4.26 + 0.14<sup>a</sup> | 4.08 + 0.21<sup>b</sup> | 4.04 + 0.13<sup>b</sup> |
| Moisture | 44.90 + 1.98<sup>b</sup> | 46.2 + 2.04<sup>b</sup> | 52.8 + 2.11<sup>a</sup> | 45.60 + 1.43<sup>b</sup> |
| Ash | 1.63 + 0.09<sup>a</sup> | 1.40 + 0.09<sup>c</sup> | 1.55 + 0.02<sup>b</sup> | 1.65 + 0.04<sup>a</sup> |
| Crude fibre | 17.90 + 1.23<sup>b</sup> | 16.08 + 1.41<sup>d</sup> | 17.02 + 1.37<sup>c</sup> | 18.40 + 1.14<sup>a</sup> |

### Phytochemical analysis

Methanolic extracts from the leaves and stems of *Celosia argentea* revealed the presence of phytochemicals, such as alkaloids, saponins, and polyphenols, such as flavonoids (Table 2a). The alkaloid, saponin, and flavonoid contents of *C. argentea*-grown aquaponics were not significantly different from those of *C. argentea* grown in the unamended loamy soil. However, these values were significantly lower than those of *C. argentea* grown in NPK-supplemented soil and in inorganic hydroponics. A similar trend was observed for *C. olitorius*, with the alkaloid, saponin, and tannin content of those grown in aquaponics similar to that of those grown in unamended loamy soil, and significantly lower than those grown in NPK-supplemented soil and inorganic hydroponics. However, there was no significant difference in the flavonoid content of *C. olitorius* grown on the four growth media (Table 2b). The four identified phytochemicals in aquaponics grown *in Ocimum gratissimum* were not significantly different from those grown in other growth media, with the exception of NPK-supplemented soil, which had significantly higher phytochemical contents (Table 2c).

**Table 2a:**
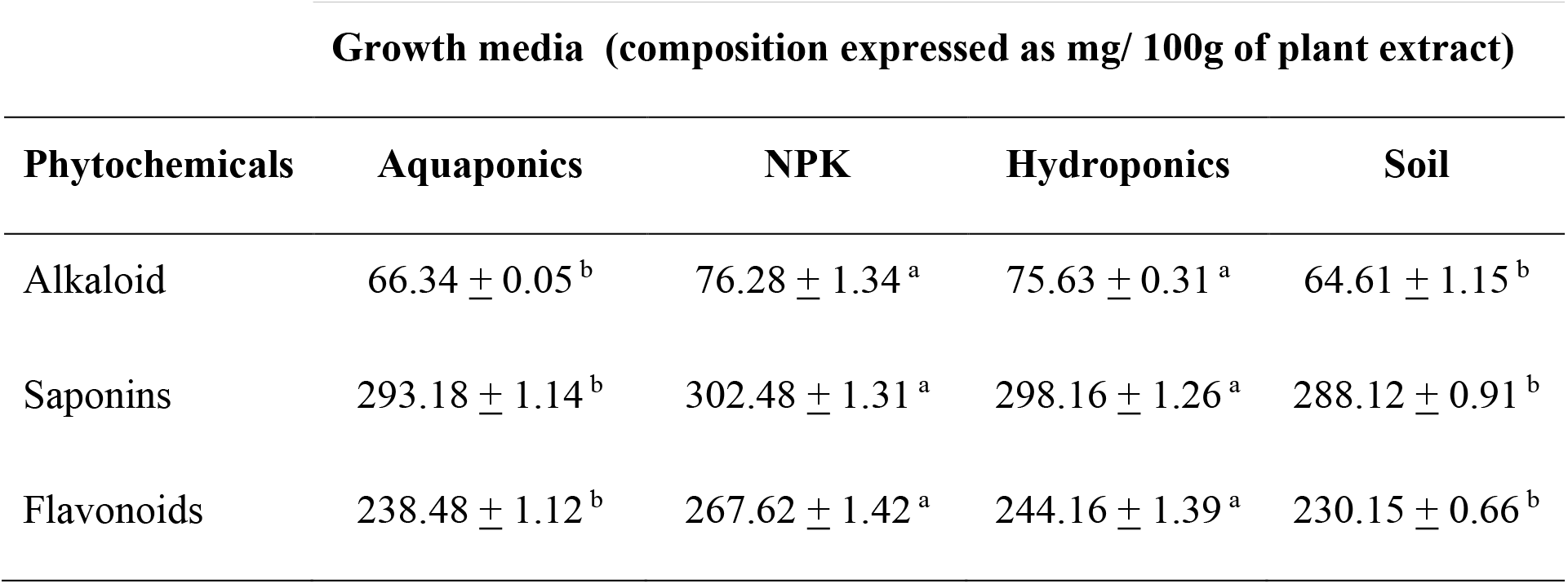
Phytochemicals in methanolic extract of *Celosia argentea*. Values with different letters are significantly different across the row at α level of 0.05.

| Growth media (composition expressed as mg/ 100g of plant extract) |  |  |  |  |
| --- | --- | --- | --- | --- |
| Phytochemicals | Aquaponics | NPK | Hydroponics | Soil |
| Alkaloid | 66.34 ± 0.05<sup>b</sup> | 76.28 ± 1.34<sup>a</sup> | 75.63 ± 0.31<sup>a</sup> | 64.61 ± 1.15<sup>b</sup> |
| Saponins | 293.18 ± 1.14<sup>b</sup> | 302.48 ± 1.31<sup>a</sup> | 298.16 ± 1.26<sup>a</sup> | 288.12 ± 0.91<sup>b</sup> |
| Flavonoids | 238.48 ± 1.12<sup>b</sup> | 267.62 ± 1.42<sup>a</sup> | 244.16 ± 1.39<sup>a</sup> | 230.15 ± 0.66<sup>b</sup> |

**Table 2b:** Phytochemicals in methanolic extract of *Corchorus olitorius*. Values with different letters are significantly different across the row at α level of 0.05.

| Growth media (composition expressed as mg/ 100g of plant extract) |  |  |  |  |
| --- | --- | --- | --- | --- |
| Phytochemicals | Aquaponic | NPK | Hydroponics | Soil |
| Alkaloid | 68.65 ± 2.05<sup>b</sup> | 79.67 ± 0.95<sup>a</sup> | 78.24 ± 1.15<sup>a</sup> | 64.32 ± 1.85<sup>b</sup> |
| Saponins | 22.79 ± 1.84<sup>b</sup> | 34.32 ± 0.44<sup>a</sup> | 31.39 ± 0.86<sup>a</sup> | 22.71 ± 0.63<sup>b</sup> |
| Tannins | 287.98 ± 0.86<sup>b</sup> | 311.41 ± 1.35<sup>a</sup> | 302.42 ± 0.92<sup>a</sup> | 272.46 ± 0.31<sup>b</sup> |
| Flavonoids | 182.92 ± 0.38<sup>a</sup> | 183.41 ± 0.34<sup>a</sup> | 182.14 ± 0.46<sup>a</sup> | 182.04 ± 0.11<sup>a</sup> |

**Table 2c:** Phytochemicals in methanolic extract of *Ocimum gratissimum*. Values with different letters are significantly different across the row at α level of 0.05.

| Growth media (composition expressed as mg/ 100g of plant extract) |  |  |  |  |
| --- | --- | --- | --- | --- |
| Phytochemicals | Aquaponic | NPK | Hydroponics | Soil |
| Alkaloid | 122.50 ± 2.05<sup>b</sup> | 134.32 ± 1.15<sup>a</sup> | 121.43 ± 0.95<sup>b</sup> | 119.41 ± 1.65<sup>b</sup> |
| Saponins | 56.78 ± 1.84<sup>b</sup> | 72.18 ± 0.21<sup>a</sup> | 49.31 ± 1.64<sup>b</sup> | 48.92 ± 1.83<sup>b</sup> |
| Tannins | 194.62 ± 0.86<sup>b</sup> | 206.11 ± 0.88<sup>a</sup> | 184.93 ± 1.12<sup>b</sup> | 189.16 ± 0.61<sup>b</sup> |
| Flavonoids | 127.8 ± 1.12<sup>b</sup> | 159.62 ± 1.32<sup>a</sup> | 124.21 ± 0.69<sup>b</sup> | 121.35 ± 1.76<sup>b</sup> |

### Test of *Corchorus olitorius* mucilage properties

The palatability of *C. olitorius* soup is determined by its viscosity (Nasir-Naeem *et al*., 2021), we therefore assessed the mucilage properties of cooked *C. olitorius* harvested from the four growth media. The viscosity of *C. olitorius* grown in aquaponics was similar to that of *C. olitorius* grown in inorganic hydroponics and in unamended loamy soil (Figure 5). However, *C. olitorius* grown in NPK-supplemented soil had the lowest viscosity as it took a longer period of time to flow.

**Figure 5:**
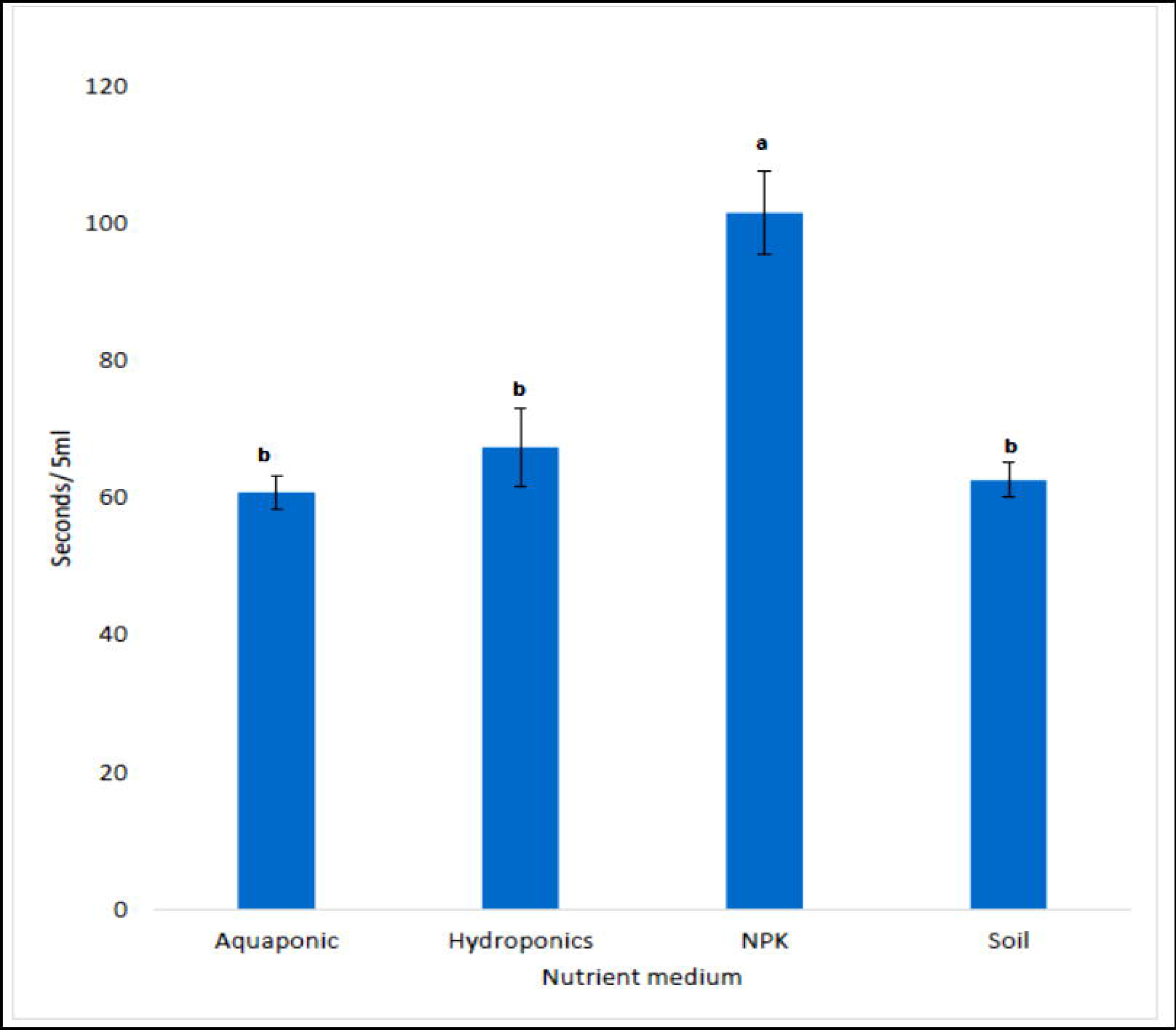
The viscosity of *Corchorus olitorius* across the 4 growth medium. Plotted means on the same horizontal axis, represented with different letters are significantly different at p= 0.05. *C. olitorius* grown in NPK supplemented soil was the least viscous while the viscosity of *C. olitorius* grown in the other growth medium were not significantly different from each other.

### Nitrogen dynamics in the aquaponics system

Efficient management of nitrogen dynamics in aquaponics is crucial for the success of the system (PETREA *et al*., 2014) as plant cultivation and aquaculture platform. Hence, this study assessed ammonia, nitrate, and nitrite concentrations at water collection points in aquaponic fish tanks, control aquaculture tanks, aquaponic recycled water reservoirs, and various plant cultivation beds. Water collected from the aquaponic fish tank was collected before it was released to the plant growth beds, whereas water collected from the control fish tank was collected prior to weekly water change. Water was collected from the plant growth beds approximately nine days after they were released from the fish tank, and water was collected from the aquaponic water reservoir prior to releasing the water back to the fish tank. This provides an insight into how efficiently each plant species manages the breakdown of ammonia from the fish tank to nitrite and nitrate. It also provides insights into the overall efficiency of aquaponic systems in managing nitrogen dynamics. The ammonia concentration (mg/l) was significantly higher in both the aquaponic fish tank and the control fish tank than at the other collection points until the 8th week (Figure 6a). In the 10th week, there was no significant difference in ammonia concentration at all collection points, except for the aquaponic recycled water reservoir (Figure 6a). The nitrite concentration was significantly higher in the control fish tank than at all other water collection points throughout the 10 weeks of observation (Figure 6b). However, the aquaponic recycled water reservoir maintained the lowest nitrite content throughout the study period. Similarly, the concentration of nitrate was significantly higher in the control fish tank than at the other water collection points. The aquaponic recycled water reservoir had the lowest nitrate concentration throughout the study period (Figure 6c). The three plant species growth beds had significantly reduced ammonia, nitrite, and nitrate concentrations compared to the aquaponic and control fish tanks.

**Figure 6:**
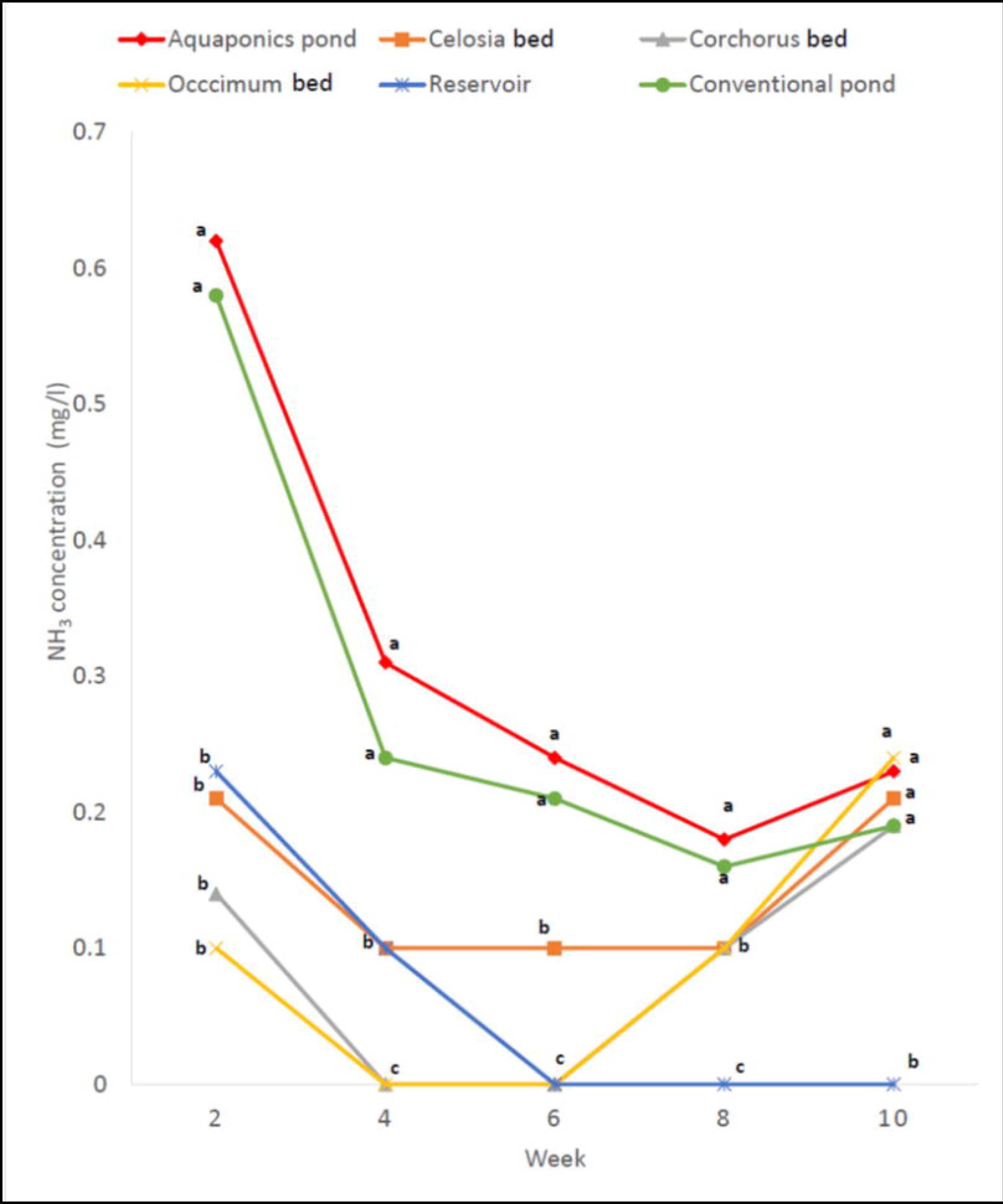
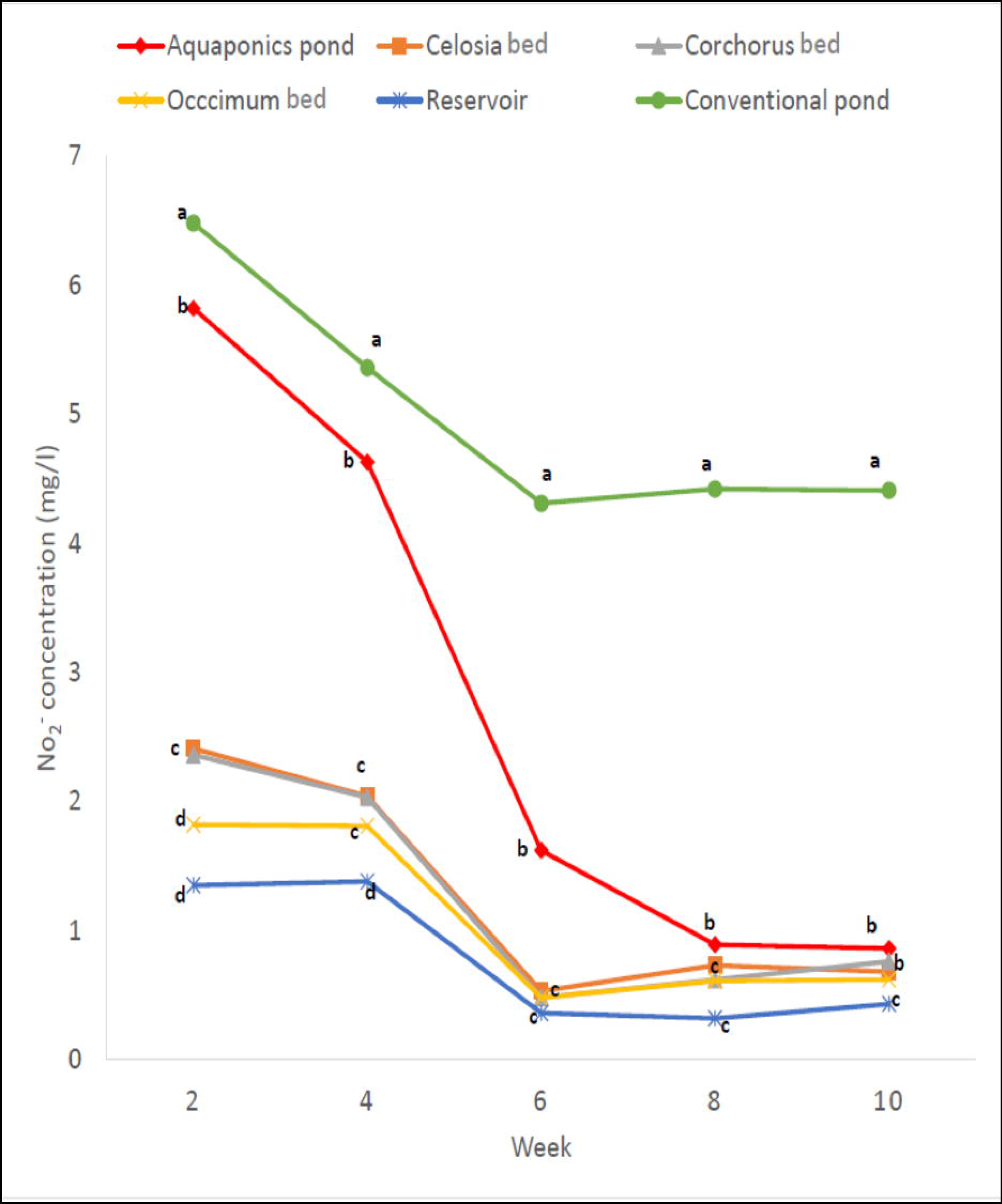
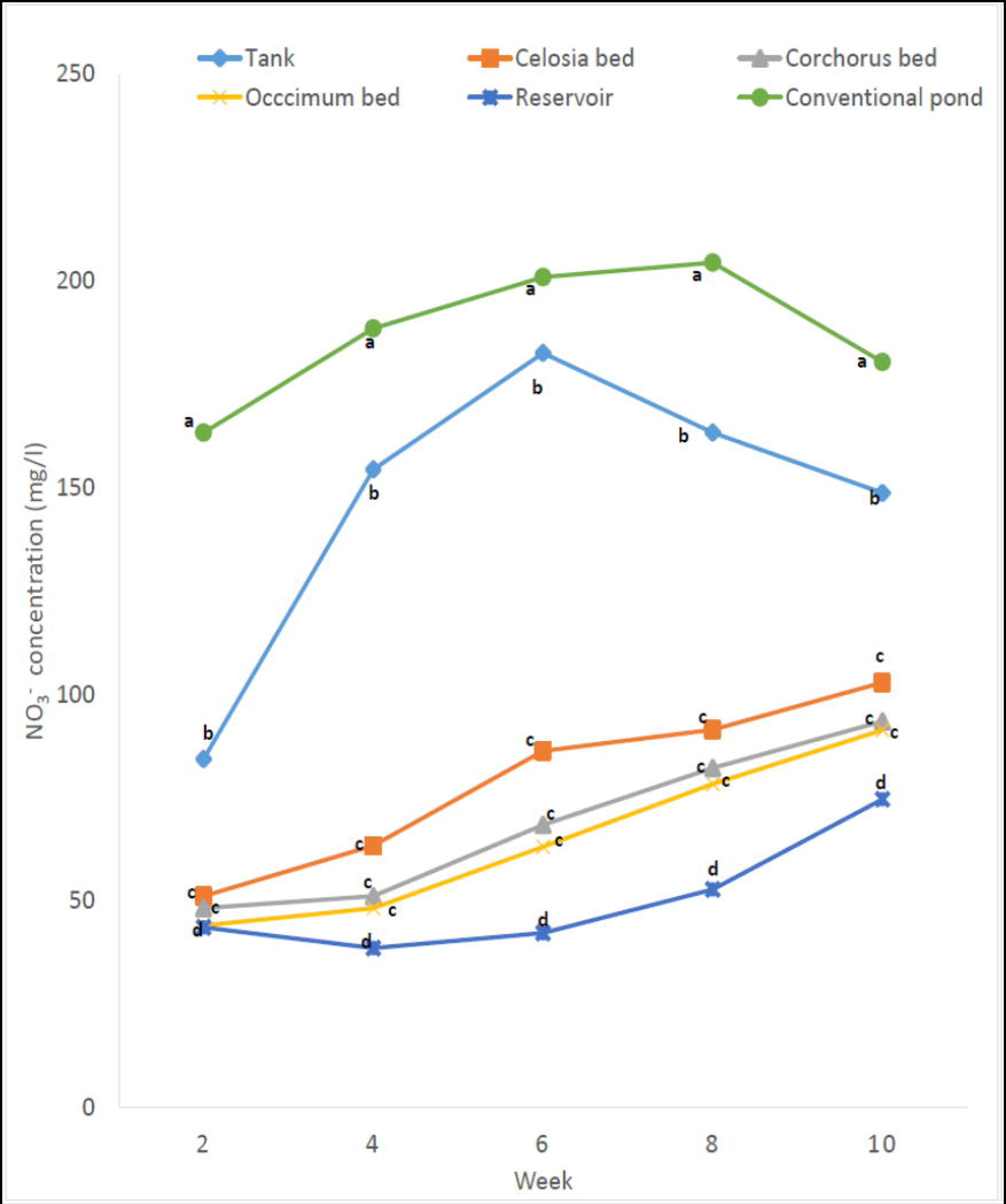
Comparison of nitrogen dynamics in aquaponics and control fish tank. **a**. Concentration of ammonia in mg/l **b**. Concentration of nitrite ion in mg/l **c**. Concentration of nitrate ion in mg/l. Aquaponic pond (red), *C. argentea* growth bed (orange), *C. olitorius* growth bed (grey), *O. gratissimum* growth bed (yellow), aquaponics water reservoir (blue) and the control fish tank (green). Plotted means, in the same week and on the same horizontal axis, represented with different letters are significantly different at p= 0.05.

## DISCUSSION

### Effects of the Aquaponics system on plant physiological growth

Environmental conditions in the aquaponic system have been reported to affect the bioavailability and uptake of nutrient elements by plants growing in the system (Roosta, 2011), influence the suppression of plant pathogenic attacks (Goddek *et al*., 2019) and consequently affect the overall physiological status of the plant. The results of this study indicate that plants grown in aquaponics have healthy growth and development comparable to those grown in unamended organic loamy soil, and better than those grown in hydroponics supplemented with inorganic mineral salts. Notably, the vegetative yield in aquaponics was less than the yield from the inorganic NPK-supplemented soil across all three plant species, whereas they all had higher biomass yields in aquaponics than in inorganic hydroponics. This might be related to defects in the management of chemical nutrient availability in the hydroponic solution (Sambo *et al*., 2019). This defect also occurs in aquaponics when there is either an inadequate or an excess amount of nitrate supplied from the aquaculture tank. However, this can be easily rectified by constant monitoring of the nitrogen cycle in the system and by prompt adjustment of the fish-to-plant ratio in the aquaponic setup (Suhl *et al*., 2016). Similarly, the significantly low vegetative productivity in inorganic hydroponics may be associated with the lack of proper aeration in the slope culture hydroponic system.

Significant differences were observed between the dry weights of the three plant species grown in aquaponics and inorganic hydroponics, despite the initial lack of significant differences in their fresh weights (Figure 2). This suggests that despite the growth medium being in the liquid state, plants grown in aquaponics efficiently converted nutrients to biomass, and development was related to cell division and elongation, whereas plants grown in inorganic hydroponics dedicated acquired resources to vacuolar enlargement from water storage and storage of water in tissues. Therefore, the water was removed when the plants were dried. This indicates that aquaponics may be a more reliable system for biomass production in vertical and indoor vegetable agriculture. The lack of significant differences in the vegetative yield in aquaponics and unamended loamy soil indicates that the cultivation of *C. argentea, C. olitorius, and O. gratissimum* in aquaponics is as productive as their cultivation in the field. Considering the risk of soil-borne infections, energy expended on weeding, and so on, the aquaponic cultivation of the three vegetables is preferred as washed granites are substituted for soil that cuts off soil-borne infections and weeds (Palm *et al*., 2018). The similarity in the vegetative adaption of the three plant species to the different growth media is in line with the report of Togawa-Urakoshi & Ueno (2022) who pointed out the similarity in the nitrogen usage efficiency, growth, and biomass partitioning of C3 plants.

### Effects of the Aquaponics system on plant nutritional and phytochemical compositions

The nutritional profiles of the three plant species revealed that vegetables grown in aquaponics had better nutritional composition than those grown in NPK-supplemented soil and inorganic hydroponics. Mofunanya *et al*. (2014) noted that the application of organic fertilizer produced *Amaranthus spinosus* with superior nutritional value compared to the application of inorganic fertilizer, while Vigar *et al*. (2019) corroborated this finding by suggesting that organic foods contain higher levels of certain nutrients and reduced pesticide content than their inorganic counterparts. The three plant species grown in unamended soil, which are sources of both organic and inorganic mineral nutrients, had increased nutritional availability compared with those grown in aquaponics. The combined application of organic manure and inorganic fertilizers has been identified as the best approach for improving crop biomass yields (Baghdadi *et al*., 2018). The results of this study also corroborate reports that *C. argentea, C. olitorius, and O. gratissimum* are rich in phytochemicals, as several studies have isolated biologically important compounds from their extracts (Alabi *et al*., 2020; Dung *et al*., 2021; Tang *et al*., 2016). Aquaponics produces plants that can supply biologically important phytochemicals to humans when consumed, and plants grown in aquaponics can compete favorably in this regard with plants grown in unamended loamy soil in terms of phytochemical compositions.

### From nitrogenous waste to nitrogenous manure

Plants in the aquaponic system depend on the uptake of nitrate as the main source of nitrogen (Hu *et al*., 2015), which plays an important role in preventing the accumulation of ammonia, nitrite, and nitrate in the aquaponic fish tank. The results of this study indicate that the three plant species were able to efficiently utilize the nitrite generated from poisonous ammonia waste produced by the aquaculture component of aquaponics to build biomass. Therefore, the water returned to the aquaponic tank contained significantly lower concentrations of ammonia, nitrate, and nitrite. This suggests that the aquaponic system was able to establish a microbiota of nitrifying bacteria and that the three plant species were able to effectively assimilate the nitrate produced. The significant difference observed in the concentration of nitrogenous ions between the plant cultivation beds and the water reservoir indicated that continuous nitrification occurred in the aquaponic reservoir. Hence, it is advisable that water passing into the reservoir should remain for approximately 24 h before recycling into the aquaponic fish tank. These three plant species are recommended for effective aquaponics owing to their good nutrient uptake. Further studies on the aquaponic microbiome will provide more insights into how the aquaponic system manages nitrogen cycling.

This physiological study revealed that the aquaponic system produces leafy vegetables whose growth, biomass productivity, nutritional contents and phytochemical composition can match those of existing conventional agricultural systems. Likewise, the study indicated that although NPK-supplemented soil produced a better biomass yield, aquaponics trump this with the advantage of environmental protection from non-biodegradable fertilizers, reduced losses due to infections by soil-borne pathogens, and the production of solely organic vegetables. Investment in more studies on plant cultivation in aquaponics will facilitate global sustainable food production while also projecting aquaponics as a prospective source of food for beyond earth explorations.

## Supporting information

Raw data

## Authors’ contributions

The authors confirm the contribution to the paper as follows:

Study conception and design: GOO & EOA

Data collection: GOO & AOA

Analysis & interpretation of results: GOO, EOA, DDS &AOA

Draft manuscript preparation: GOO & DDS

All authors reviewed and approved the final version of the manuscript submitted

## Acknowledgment

Thanks to Mrs. Suberu of the University of Lagos Fishery department who helped with regular feeding of the fish and her professional advice on fishery management are invaluable.

## Data access statement

All relevant data are within the paper and its supporting information files

## Ethics approval

This study was approved by the Ethics Committee of the University of Lagos, Nigeria.

